# Fss1, a CTNS/PQ domain protein, has dual roles in promoting fertility in *Schizosaccharomyces pombe*

**DOI:** 10.64898/2026.10.26.761791

**Authors:** Andrew J. Bestul, Jeffrey J. Lange, Stephanie H. Nowotarski, William B. Redwine, Sarah E. Zanders

## Abstract

Sexual reproduction is critical in most eukaryotes, but many genes required for the process remain to be identified and characterized. A previous saturation mutagenesis study in *Schizosaccharomyces pombe* implicated several unnamed genes in having roles in sexual reproduction. In this work, we provide an initial characterization of 13 of these genes and an in-depth characterization of one gene, *fss1* (facilitates <u>s</u>tarvation <u>s</u>ex). Fss1 has high structural homology to the human PQ-loop repeat family PQLC2 transceptor (amino acid transporter plus membrane receptor), but likely lacks the ability to transport amino acids due to loss of a critical PQ motif. *fss1Δ* mutants have no defects in meiosis but have pronounced defects in packaging the products of meiosis into spores, particularly in nutrient-poor environments. The spore formation phenotypes are recessive as *fss1Δ* spores produced by a *fss1+*/ *fss1Δ* heterozygote develop normally. Immuno-gold electron microscopy shows Fss1 localizes to vesicles that fuse to the ephemeral outer spore membrane as the spore wall matures below. Fss1 protein is also produced within spores and this population is required for normal vacuole morphology and timely germination. We propose a model in which Fss1 is a vesicle receptor that facilitates spore wall formation from diploid-expressed protein, while spore-expressed protein acts at the vacuole to promote an internal cellular environment that facilitates germination. This work expands our catalog of genes that contribute to sexual reproduction and identifies two previously unknown tasks in spore biology that are both facilitated by Fss1.

## Introduction

Most eukaryotes benefit from sexual reproduction. In some, sex is not essential, but it still plays critical roles in facilitating adaptation and long-term evolutionary success. In other organisms, like humans, sex is the sole form of reproduction. Despite the ubiquity and importance of sex, many facets of the process are not well-understood at the molecular level.

Sexual reproduction requires meiosis to generate gametes from diploid progenitors and subsequently requires two gametes to fuse to regenerate a novel diploid. Much of what we know about the molecular mechanisms of sexual reproduction was first discovered in simple model plants, fungi and insects [1–3]. These foundational studies in model systems were critical as many aspects of meiosis, gamete development and gamete fusion are broadly conserved, including in humans [4]. Still, sexual reproduction remains incompletely characterized in even the simplest species. This indicates that model systems have the potential to identify additional, potentially conserved, molecular mechanisms supporting sexual reproduction [5–11].

The fission yeast *Schizosaccharomyces pombe* is a long-established, highly tractable model system for understanding sexual reproduction [12, 13]. *S. pombe* cells are generally haploid and can multiply via mitosis when nutrients are abundant. The cells can be one of two mating-types, but as the cells divide, they switch between the two types such that a clonal population contains both mating-types. If the cells become starved for nitrogen, cells of opposite mating types can fuse to make a diploid zygote. Under persistent nitrogen starvation, this zygote undergoes meiosis to generate four haploid cells [14, 15]. These haploid cells undergo a developmental process known as sporulation in which the four meiotic products are first enveloped in a double forespore membrane via a vesicle fusion process [16, 17]. A thick spore wall then develops between the two cell membranes, and the outer membrane is eventually lost. The result is four long-lived, stress resistant [18], dormant spores packaged together in an ascus. Spores can germinate and resume mitotic growth when returned to a nutrient-rich environment [16, 19].

Many key contributors to sexual reproduction have been identified by screens in *S. pombe*, or due to homology to factors identified in other model systems, including the budding yeast *Saccharomyces cerevisiae* [5, 11, 20–24]. Still, classic screens in both organisms were generally not saturating and are best at detecting severe phenotypes. To get a wider view of the genes that contribute to sexual reproduction, our group previously executed a genome-wide saturation mutagenesis to quantify the contribution of non-essential genes to the production of viable spores in *S. pombe*. That work implicated 142 additional genes as unannotated contributors to sexual reproduction [25].

Here, we follow up on that saturation mutagenesis to individually assay how 14 candidate genes contribute to sexual reproduction. We chose genes that were unnamed at the outset of our study to maximize our chances of finding novel phenotypes and to further efforts to characterize all *S. pombe* genes [26, 27]. We assayed each mutant for mating efficiency, sporulation efficiency and spore germination. Each mutant exhibited a significant defect, ranging from mild to moderate severity, in one or more aspects of sexual reproduction.

We chose one gene, *SPAC4C5.03*, for further characterization due to the relative severity and breadth of the phenotypes observed upon initial characterization. We named this gene, *fss1*, for **f**acilitates **<u>s</u>**tarvation **<u>s</u>**ex, as the mutant phenotypes associated with loss of the gene are exacerbated on low nutrient sporulation media. Fss1 does not have clear sequence orthologs outside of fungi but shares structural homology to human PQLC2, a transceptor protein that acts as an amino acid transporter and receptor on lysosomes [28–30]. Members of the same protein family (CTNS/PQ-loop domain proteins; MtN3/SLV/SWEET proteins in plants) have previously been implicated in promoting fertility in plants [31, 32].

In *S. pombe,* we find that Fss1 is required in diploids to promote spore wall development. Our experiments also illustrate a novel phase of spore wall development where Fss1-bearing vesicles are deposited to the short-lived outer spore membrane as the spore wall develops. Within haploid spores, Fss1 is required to establish normal vacuole morphology and to expedite germination. Altogether, this work expands our understanding of genes and molecular processes required for sexual reproduction in *S. pombe* and can potentially guide studies of homologous genes and mechanisms in other eukaryotes.

## Results

### Survey of 14 candidate sexual reproduction genes, including identification of <u>s</u>ocial <u>d</u>istancing <u>g</u>ene 2 (*sdg2*)

Billmyre et al used saturating transposon mutagenesis paired with sequencing (TN-Seq) to implicate over 100 novel genes as contributing to sexual reproduction in *S. pombe*, including over 40 genes that lacked an assigned name at the time of that study [25]. Briefly, the TN-seq experiment assayed the frequency of transposon insertions in each gene in a large, diverse library both before and after sexual reproduction (spore formation). Transposon insertions in genes that promote sexual reproduction were depleted from the library after sex. Here, we choose 14 of those largely uncharacterized genes with strong depletion in mutations after sex for characterization (Table S1). Eight of these genes have human orthologs, or share significant structural homology with a human protein (according to AlphaFold prediction), five genes appear to be fungal specific, and one gene (*SPAC24C6.04, put2*) is orthologous to genes found in bacteria, archaea and humans (Table S1) [26].

We generated complete coding sequence deletion mutations of the 14 candidate genes and first assayed if they exhibited one of the two known false positive phenotypes identified in Billmyre et al. that erroneously led genes to be implicated in sexual reproduction: 1) slow growth or 2) a density dependent growth phenotype defined by the **<u>s</u>**ocial **<u>d</u>**istancing **<u>g</u>**ene, *sdg1* [25]. None of the mutants exhibited markedly slow growth compared to wild-type on solid rich media (YEA+Sup; Figure S1A). Deletion of *SPBC1604.03c* did, however, lead to a density-dependent growth phenotype like that of *sdg1Δ* mutants (Figure S2A). Specifically, the *SPBC1604.03c* mutants often failed to grow when plated at low cell density, but this phenotype could be at least partially rescued by growing the cells on conditioned media (Figure S2B-D). Based on this phenotype, we named *SPBC1604.03c sdg2* (<u>s</u>ocial <u>d</u>istancing <u>g</u>ene 2) and suspected the mutant’s inability to grow at low cell density at least partially contributed to the gene’s identification in our TN-seq assay.

We tested each mutant’s ability to complete sexual reproduction by plating cultured haploid cells onto media that induces mating and spore formation. We then imaged the cells and assayed the mating efficiency (percent mated cells), sporulation efficiency (percent of zygotes that form at least one spore), and the fraction of atypical asci produced by each mutant. We also isolated spores produced by each mutant and assayed their ability to germinate (germination rate) and the time required for germination (Figure S1B-D; Table S1). Finally, we measured the ‘viable spore yield’ of each mutant as a coarse, aggregated quantification of overall fertility. The viable spore yield assay quantifies the number of viable spores produced per viable cell placed on sporulation medium (Figure 1A). For these experiments, we used two types of synthetic sporulation media: SPA and SPAS, which is SPA supplemented with additional nucleotides and amino acids (See methods) (Figure S1E). We elected to not use the non-synthetic MEA sporulation media used in the original TN-seq experiment as we obtain more consistent results with the synthetic media.

**Figure 1:**
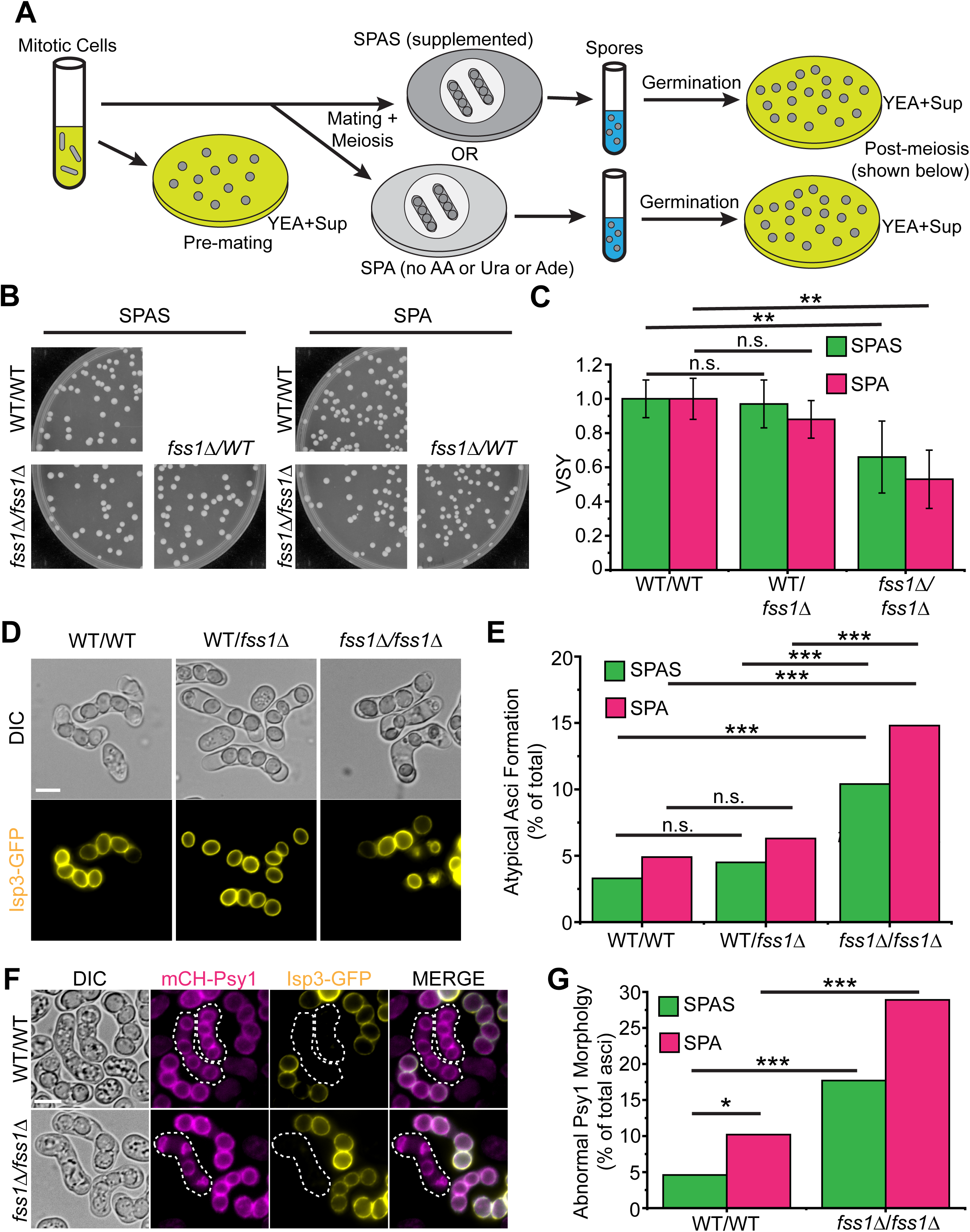
*fss1Δ* confers recessive defects in viable spore yield and spore formation. A) Diagram illustrating the viable spore yield (VSY) assay using two different mating media: SPAS (supplemented with leucine, uracil, lysine, histidine and adenine) and SPA (no additional supplements). Colony forming units (CFU) on rich media (YEA+Sup) are calculated before (mitotic cells) and after sexual reproduction (spores) and the VSY is calculated as the (post sex CFU/pre sex CFU) (See Materials and Methods for specific protocol). B) Example images of the YEA+Sup spore germination plates from spores formed on SPAS media (left) or SPA media (right) by the indicated diploids. C) The average VSY, normalized to wild-type, is shown for the indicated diploids on both mating media. Each diploid was assayed six times in both media. n.s.= not significant; **p<0.001; t-test. D) Asci produced by the indicated diploids after 24 hours on SPAS. Isp3-GFP marks the spore exterior. Scale Bar, 5 µm. E) Percentage of atypical asci, assayed from Isp3-GFP DIC images (as in D), produced by the indicated diploids on both mating media. Additional examples of atypical asci are shown in Figure S1C. n>200 for all strains; n.s.= not significant; ***p<0.0001, chi-squared test. F) Asci produced by the indicated diploids after 16 hours on SPAS. mCherry-Psy1 marks the forespore membrane which forms prior to the spore wall, marked by Isp3-GFP. Scale bar, 5 µm. G) Percentage of asci produced by the indicated diploids with atypical prospore membrane morphology (i.e. mCherry-Psy1 in F) prior to Isp3-GFP deposition. Asci at the targeted stage are outlined with a dotted line in F. Values for asci produced on both types of mating media are shown. n>50 for all strains; *p<0.05; ***p<0.0001, chi-squared test.

Overall, more mutant phenotypes were observed on the more stringent sporulation media, SPA, so we will briefly describe those results in the text, although all our complete results are summarized in Table S1. *sdg2Δ* mutants did not show significant defects in mating but did show minor defects in spore formation (Table S1). *sdg2Δ* mutants also moderately reduced viable spore yields, analogous to *sdg1Δ* mutants tested in parallel. We cannot distinguish if these reproductive phenotypes are due to incomplete suppression of the density dependent growth phenotype on the media used for these experiments or due to separate roles for these genes in sexual reproduction, although we suspect the former, due to our previous observations of *sdg1Δ* mutants [25].

The other 13 genes all showed a significant defect in one or more assay (Table S1). The most common defect, shared by 11 mutants, was a loss of viable spore yield from mating on the minimal SPA media. Deletion mutants of two genes, *SPBC530.11c* and *SPAC17A2.02c* (now known as *tlc4*), both exhibited reduced mating efficiency on SPA media. Four of the hits (*SPAC4C5.03, dcw1, sdg2 and ank1)* showed increases in atypical asci formation. *SPAC4C5.03* mutants consistently showed the highest rate of spore formation defects and a large loss of viable spore yield in SPA. These phenotypes led us to believe the gene was responsible for facilitating meiosis and spore formation in harsh conditions, so we named the gene, <u>F</u>acilitates <u>S</u>tarvation <u>S</u>ex (*fss1*) and pursued further characterization of the gene.

### Fss1 is a predicted PQ-loop repeat family protein

The *fss1* gene is broadly conserved in fungi, but is not found in the budding yeast *Saccharomyces cerevisiae.* Fss1 is one of three predicted members of the PQ-loop repeat family of transmembrane proteins in *S. pombe*, along with the characterized Stm1, a vacuolar amino acid transporter and the uncharacterized SPAC2E12.03c (Figure S3A) [33, 34]. PQ-loop proteins have a pair of PQ repeats on two spanning transmembrane helices connected by a loop. These two PQ domains provide flexible hinges that enable a conformational change that allows substrate transfer across membranes [35, 36]. The Fss1 AlphaFold structural prediction (Figure S3B) is moderately similar to *S. pombe* Stm1 and human PQLC2, but resembles human cystinosin (CTNS), another PQ-loop protein, the most structurally.

The site of the first PQ in Fss1 is, however, an LQ in Fss1 and its *Schizosaccharomyces* orthologs (Figure S3A). This mutation is predicted to disrupt the hinge required for transport and this same mutation (P to L in the first PQ domain) strongly disrupts amino acid transport in the *C. elegans* ortholog of PQLC2 [35]. Molecular structural predictions support that Fss1 is likely to be similarly unable to complete the conformational changes required for transport (Figure S3C). The Fss1 structural prediction also has an extended C-terminal helix not seen in other PQ-loop repeat proteins in *S. pombe* (Figure S3B), which we speculate could contribute to a membrane receptor function [37]. This C-terminal region contains a site for TORC1 phosphorylation (S288) during nitrogen starvation and TORC1 activity promotes sexual reproduction [38].

### Fss1 promotes sexual reproduction, particularly in nutrient-depleted conditions

The original screening of the *fss1Δ* phenotype was done in a *leu1, ura4* auxotrophic strain background (Table S1). The *leu1* auxotrophy has been linked to mating and fusion deficiencies in *S. pombe* and leucine promotes TORC1 activity [38, 39]. Given Fss1’s more severe mutant phenotype on stringent sporulation media, we suspected the mutant phenotype might be dependent on auxotrophies. We tested this by assaying viable spore yield in a prototrophic background. We found that the *fss1Δ* mutant viable spore yield phenotype was not completely dependent on the auxotrophies as the prototrophic *fss1Δ* mutant also had reduced viable spore yield on both SPA and SPAS media (Figure 1B, C; p<0.005, t-test). This phenotype was recessive, as we found no significant reduction in viable spore yield of *fss1Δ* /*fss1*+ heterozygotes (Figure 1B, C; p>0.05, t-test).

We next tested if the *fss1Δ* phenotypes were exacerbated by auxotrophies by comparing the viable spore yields of wild-type and *leu1* auxotrophs. The *leu1-32* mutation reduced viable spore yield in both wild-type and *fss1Δ* mutant backgrounds. The *leu1-32* mutation reduced wild-type viable spore yield by about 30%, while it reduced viable spore yield of *fss1Δ* mutants by 50-60%. However, this difference was not statistically significant (Figure S4A, B; p>0.05, chi-squared test). Together, our experiments demonstrate that Fss1 promotes sexual reproduction and is particularly important when cellular nutrients are low.

### *fss1Δ* diploids exhibit defects in spore formation

To assay where the *fss1Δ* mutant defects occur, we used a tagged histone marker (*hht1-mRFP)* to visualize nuclei through sexual reproduction on SPAS media. We were unable to complete these experiments on the more stringent SPA media, as even the wild-type cells struggled to complete meiosis under our imaging conditions. We observed no defects in mating (cell fusion) or karyogamy (nuclear fusion) in *fss1Δ* mutants. The *fss1Δ* mutants also progressed through meiosis similar to wild-type with a non-significant increase in cells that fail to complete MII (Figure S5A-C; p>0.05, chi-squared test). We also tested chromosome segregation fidelity in the first meiotic division by assaying the rates of disomy for chromosome 3 amongst viable spores produced on both SPA and SPAs (Figure S5D, E). We saw no defects relative to wild-type. These data suggests that the defects in *fss1Δ* mutants arise after the meiotic divisions.

To assess spore formation, we used Isp3-GFP, which marks the outer spore wall [40] to visualize spore morphology in wild-type and *fss1Δ* prototrophic strains (Figure 1D). Asci created by a *fss1Δ* homozygous diploid were more likely to be atypical in that they contained spores with abnormal appearance or less than four spores (Figure 1E; p<0.0001, chi-squared test). As with the viable spore yield phenotype, this defect was most penetrant when the tetrads were produced on the stringent SPA medium (Figure 1E). This spore formation phenotype is recessive, as we observed wild-type levels of atypical asci in *fss1Δ/fss1+* heterozygotes (Figure 1D, E). To test how early *fss1Δ* diploids fail in spore formation, we looked at mCherry-Psy1, a marker for the forespore membrane prior to the deposition of Isp3 (asci outlined with dotted lines in Figure 1F). Asci produced by *fss1Δ* diploids generally contained four areas of Psy1 signal, but these exhibited abnormal morphology at much higher rates than wild-type (Figure 1F, G; p<0.0001, chi-squared test), Thus, our results demonstrate that Fss1 protein is important in the diploid stage to support early spore formation.

### The Fss1 protein localizes to cytoplasmic vesicles during meiosis and then to the spore periphery upon spore formation

To determine where the Fss1 protein localizes during sexual reproduction, we generated an ectopic GFP-tagged allele under the control of the *41nmt1* promoter, *41nmt1-GFP-fss1*. We integrated this allele into the genome at its native locus and found that the allele largely phenocopies the wild-type allele for ascus morphology, but has a viable spore yield on SPA that is intermediate between those of wild-type and the *fss1Δ* mutant (Figure S6A-D). These data indicate that the allele is functional, but not fully wild-type. We imaged cells containing both the *41nmt1-GFP-fss1* allele and the tagged histone, *hht1-mRFP*, as they progressed through sexual reproduction. We could first detect GFP soon after karyogamy and it was dispersed throughout the cytoplasm. At the end of the second meiotic division, GFP-Fss1 localized at the spore perimeter and gained fluorescence intensity as the spore matured (Figure 2A).

**Figure 2:**
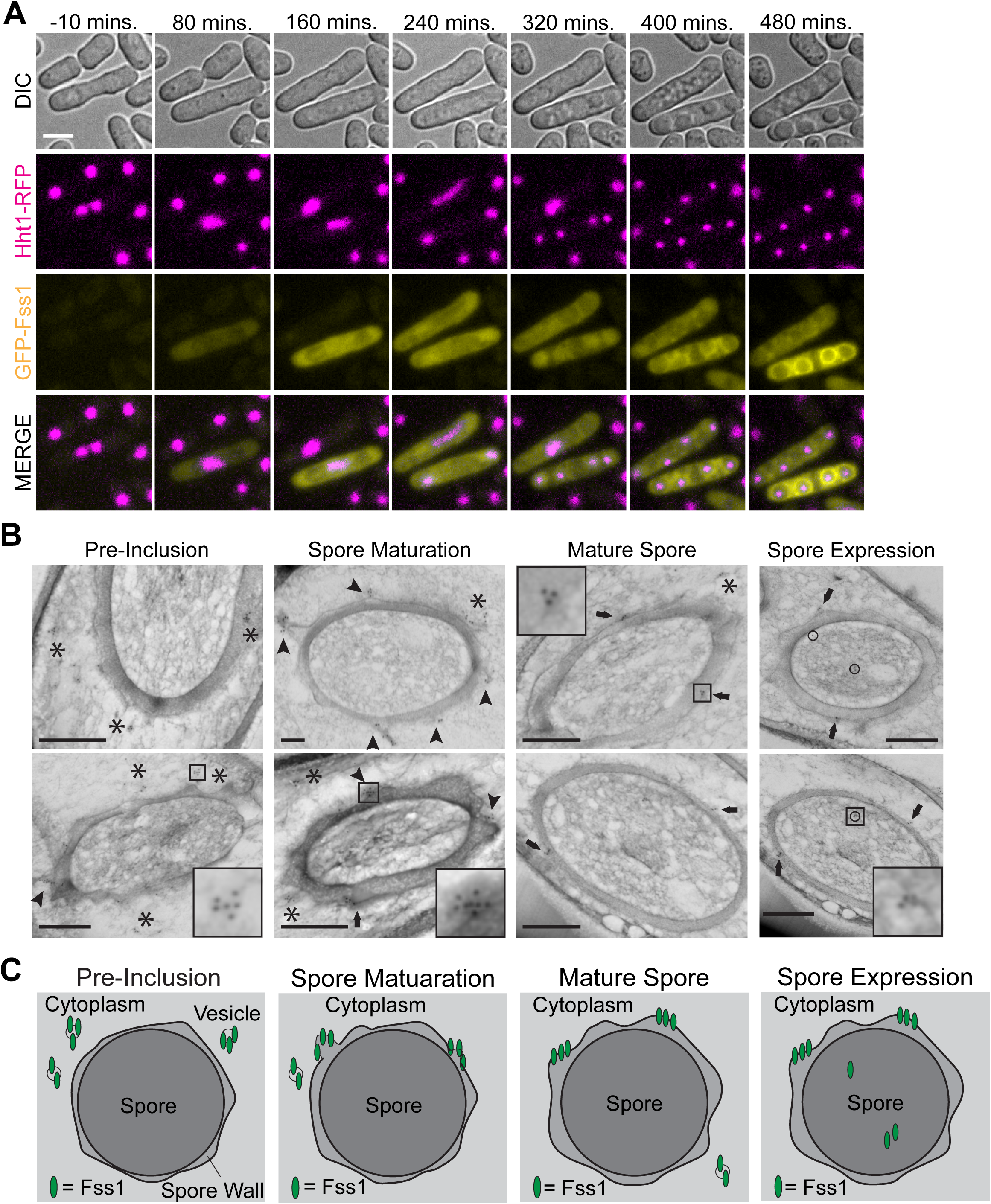
Fss1 localizes to vesicles during meiosis and then fuses into the spore wall during spore maturation. A) Timelapse images of *41nmt1-GFP-fss1* homozygous diploids undergoing meiosis on SPAS plates with Hhht1-RFP marking the nucleus. The zero timepoint (t=0, not shown) is karyogamy. Scale bar, 5 µm. B) Images of TEM slices during spore formation and maturation with GFP-Fss1 immuno-labeled with nano-gold particles. The cells were staged by Fss1 localization. Fss1 starts in the epiplasm surrounding the spores (pre-inclusion; marked with asterisks) then Fss1-bearing vesicles fuse to the outer spore membrane during spore maturation (marked with arrowheads), where the protein remains in mature spores (marked with arrows). Additional Fss1 localizes within the cytoplasm of mature spores (circled). Boxes indicate insets to better illustrate the clumped nano-gold particles for each type of localization. Scale bars, 500 nm. C) Cartoon depicting localization of Fss1 during spore formation and maturation, going from epiplasmic (Pre-inclusion) to fusing vesicle (Spore Maturation) to integrated with the spore wall (Mature Spores) to localization in mature spores (Spore Expression).

We found the initial diffuse localization of GFP-Fss1 surprising as the predicted protein structure has seven transmembrane domains (Figure S3B, C). To probe this localization pattern further, we performed immunoelectron microscopy to visualize GFP-Fss1 in an unsynchronized population of cells undergoing sexual reproduction (Figure 2B). We observed the GFP-Fss1 localized in four distinct patterns during spore formation and maturation: 1) cytoplasmic clusters of 2-4 gold particles (asterisk); 2) clumps of gold particles associated with an electro-dense mass that appears to be fusing with the outer spore membrane (arrowhead); 3) clusters of gold particles that are incorporated into the outer regions of the spore (arrows); and 4) gold particles inside mature spores (circles; Figure 2B). These localizations were specific to GFP-Fss1, as just using the secondary antibody or cells with untagged Fss1 showed very few gold particles, including zero at the localization sites shown in Figure 2B (Figure S7).

We infer from these localization patterns that the initial diffuse fluorescence signal and the small cytoplasmic clusters represent small GFP-Fss1 containing vesicles. We were unable to clearly visualize Fss1-bearing vesicle membranes prior to fusion with the spore wall, but our inference at this stage is based on the transmembrane domains of Fss1 and the clustered nature of the immunogold particles prior to assimilation into the developing spore wall. The vesicle inference has stronger support during spore maturation as our images show the immunogold particles joining the spore wall via a vesicle fusion process with the outer spore membrane. The immunogold signal remains on the outer spore wall through the rest of spore maturation. It is well-established that a process of vesicle fusion builds the initial two layers of the forespore membrane [41]. Our images reveal that a second wave of vesicle fusion occurs after the forespore membrane closes. These vesicles fuse to the outer spore membrane as the spore wall develops below (Figure 2B, C). We also considered our immunogold images could reflect blebbing out from the outer spore membrane but ruled this out for two reasons: 1) the progressive increase in GFP signal at spore peripheries shown in Figure 2A and 2) the expansion of the outer spore membrane as the spore wall grows would require additional membrane. We suspect that the Fss1 signal within spores is a separate population produced within the spores, and we discuss this population in a later section.

### Fss1 is needed in the diploid to form a proper spore wall during spore maturation

To better understand how Fss1 supports spore formation, we used scanning transmission electron microscopy (STEM) to look at cross sections of spores produced by wild-type and *fss1Δ* mutant diploids on both SPA and SPAS. We suspect that some of the most abnormal spores (e.g. Figure 1F) produced by the *fss1Δ* diploids were lost during the processing steps that proceeded imaging (see methods), but differences from spores produced by wild-type diploids were still apparent. We found that the surface of the spores produced by *fss1Δ* diploids sometimes appeared smoother than those produced by wild-type (Figure 3A) and the spore walls were significantly thinner (Figure 3B; p<0.0001; t-test).

**Figure 3:**
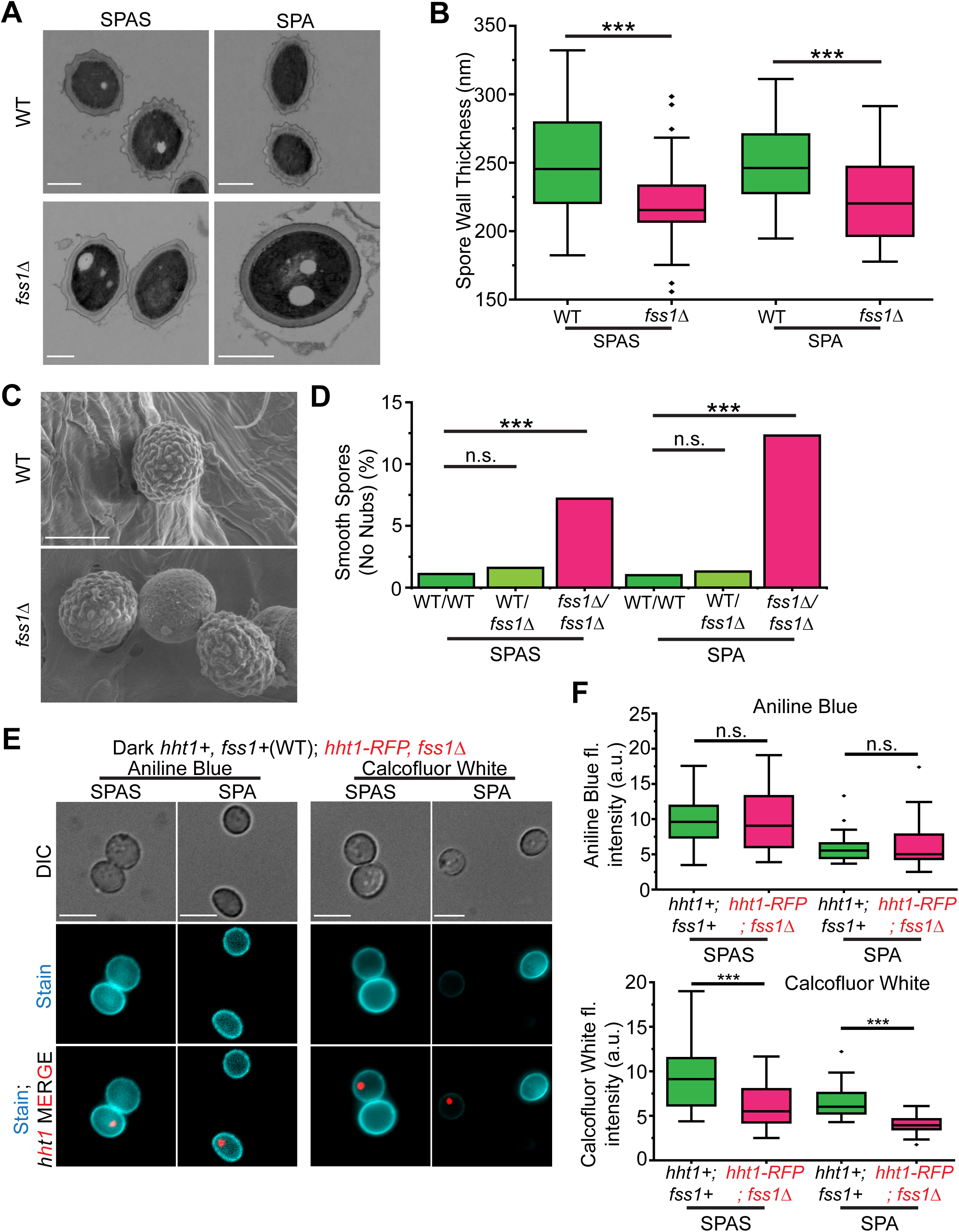
Fss1 expression in diploids promotes proper spore wall formation. A) STEM images of cross-sections spores of the indicated genotypes produced by homozygous diploids on SPAS or SPA. The electrodense dark center is the spore cytoplasm, while the lighter wavy layer surrounding the center is spore wall. Scale bars, 1000 nm. B) The thickness of the spore wall from STEM images. n>30 for all samples; ***, p<0.0001, t-test. C) SEM images of the surfaces of mature spores produced on SPAS by diploids of the indicated genotypes. Scale bars, 1000 nm. D) The percentage of spores of the indicated genotypes produced on either SPAS or SPA that had smooth spore walls. n>400 for all samples; All p-values are compared to homozygous wild-type (WT/WT); n.s. = not significant; ***p<0.0001; chi-squared test. E) Images of spores produced by wild-type diploids mixed with spores produced by *fss1Δ* homozygous diploids with Hht1*-*mRFP marking the nucleus. Spores of the two genotypes were independently created on both SPA and SPAS plates and were then mixed (SPA with SPA, etc) prior to staining with Aniline Blue (left columns) or Calcofluor White (right columns) and imaging. Scale bars, 5 µm. F) The quantification of Aniline Blue (top) and Calcofluor White (bottom) fluorescence intensity of spores from the experiment shown in E. n>30 for all samples. n.s. = not significant; ***p<0.0001; t-test.

To visualize the surface of the spores, we used scanning electron microscopy. While most spores we could image that were produced by *fss1Δ* diploids looked grossly normal, we found a significant increase in the number of spores that lacked the rough surface ‘nubs’ characteristic of *S. pombe* spores. Like the viable spore yield and the fraction of atypical tetrads, this nub-less phenotype was most pronounced on the stringent SPA medium (p<0.0001; chi-squared test). Also like those assays, this phenotype was recessive in that *fss1Δ*/*fss1+* diploids were wild-type (Figure 3C, D). This indicates that these spore development phenotypes depend on the diploid genotype, not the genotype of the developing spore.

Given the altered appearance of spores produced by *fss1Δ* diploids, we tested if their molecular composition was also altered. *S. pombe* spore walls contain α-glucans, β-glucans, galactomannans and chitin, with an exterior layer of Isp3 protein [16, 42, 43]. As described above, we found that Isp3 was deposited on spores produced by *fss1Δ* diploids, sometimes even on small poorly formed spores (Figure 1D, E). To assay the β-glucans and chitin, we stained the spores with Aniline Blue (stains β-1-3-glucans) and separately, with Calcofluor White (stains mostly chitin with some staining of β-1-3-glucans) [44, 45]. To mitigate any effects from staining different batches, we stained spores generated by wild-type and *fss1Δ* diploids together and imaged them at the same time. To delineate between the two types of spores, the *fss1Δ* diploid carried the *hht1-mRFP* marker, so the spores produced by the *fss1Δ* diploid had fluorescent nuclei. We also did the reverse experiment (wild-type nuclei marked with *hht1-mRFP)* and obtained similar results (Figure S8A, B). We found no difference in Aniline Blue staining between spores produced by wild-type and *fss1Δ* diploids, implying that the β-1-3-glucan component is unaffected by loss of *fss1* (Figure 3E, F). However, spores produced by *fss1Δ* diploids show a marked reduction in Calcofluor White staining, particularly in spores created on SPA media (Figure 3E, F; p<0.0001, t-test). This indicates that Fss1 facilitates the incorporation of chitin into the spore wall.

We next wanted to test if the spore chitin assimilation defect was a phenotype of *fss1Δ* homozygous mutant diploids or of *fss1Δ* mutant spores. We tested this by staining *fss1Δ/fss1+* heterozygotes. If the phenotype was dependent on the spore genotype, we predicted the spores produced by heterozygotes should have a mix of bright and dimly stained spores, similar to an artificial mix of spores produced by wild-type and homozygous mutant diploids. Our experiment revealed no difference in variability of staining in heterozygotes compared to the variability observed in spores produced by wild-type (Figure S8C, D). These data show that the defect in incorporating chitin into spores is a phenotype of *fss1Δ* homozygous mutant diploids, not of *fss1Δ* mutant spores.

We suspected the altered morphology and spore wall composition likely caused reduced spore viability, which would explain the reduction in viable spore yield of *fss1Δ* mutants. To test this directly we assayed the ability of spores to germinate using time-lapse microscopy. As with the electron microscopy described above, we suspect that some of the most malformed spores produced by *fss1Δ* diploids were lost during the processing steps that proceeded imaging. Of the spores we could image, however, we found that those produced by *fss1Δ* diploids were significantly less likely to germinate after 16 hours on rich media (YEA+Sup). Like the viable spore yield, aberrant tetrad frequency and the nub-less phenotype, the failure to germinate was enhanced on SPA media and was recessive: spores produced by *fss1Δ/fss1+* heterozygotes germinated at a frequency indistinguishable from those produced by wild-type diploids (Figure 4A-D). Altogether, our experiments suggest the vesicle-born Fss1 protein population found in the shared cytoplasm of the developing spores promotes the development of spore walls, particularly the accumulation of chitin, on all spores in a tetrad. In the absence of Fss1, the spore development is suboptimal and spore viability is reduced.

**Figure 4:**
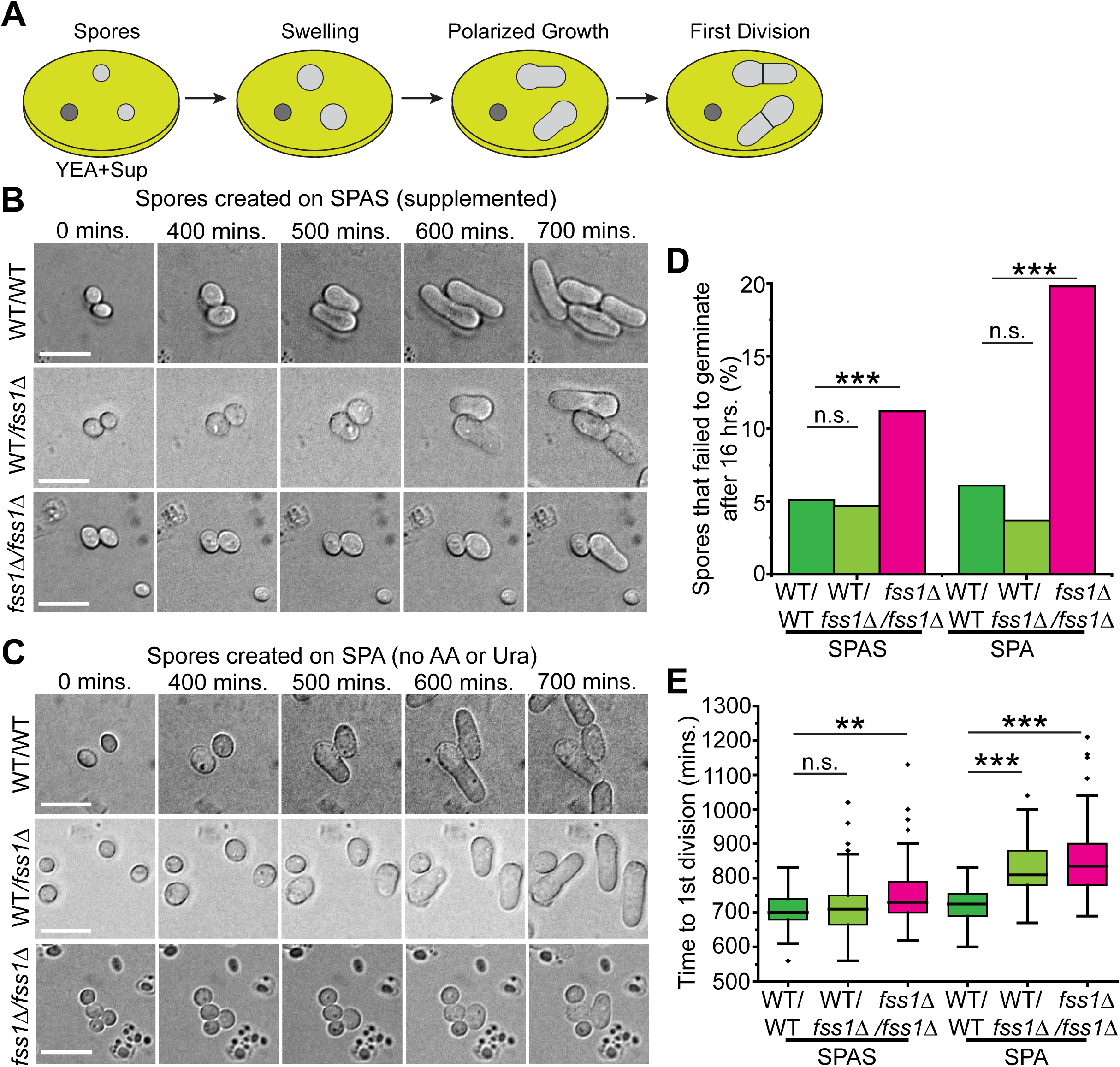
*fss1*Δ has a recessive decrease in spore germination efficiency, but spores produced by both *fss1*Δ heterozygotes and homozygotes can show delayed germination. A) Diagram showing the steps of spore germination. B and C) Timelapse DIC images of spores produced on SPAS (A) or on SPA (B) by diploids of the given genotypes germinating on YEA+Sup plates. The spores were imaged every 10 minutes for 16 hours. Scale bars, 5 µm. D) The percentage of spores from the experiments in B and C that failed to germinate within 16 hours. n>200 for all samples; n.s. = not significant; ***, p<0.0001, chi-squared test. E) First mitotic division timing for spores from the experiments in B and C. n>60 for all samples; n.s. = not significant; **p<0.001; ***p<0.0001; t-test.

### *fss1Δ* mutant spores have altered vacuole morphology and delayed germination

Although the effect of Fss1 on a spore’s propensity to germinate was recessive, our spore germination experiments discovered an additional phenotype that is not. The germination of spores produced by *fss1Δ* homozygous diploids is delayed relative to wild-type and *fss1Δ/fss1+* heterozygotes exhibit an intermediate phenotype (Figure 4E; p=0.0012, SPAS; p<0.0001, SPA; t-test). This phenotype, along with the population of GFP-Fss1 we detected within spores (Figure 2B-C), suggested that Fss1 could have a separate role within spores that facilitates timely germination.

Fss1 shares structural similarity to human PQL2 and cystinosin, which localizes to lysosomes [35, 46–48], the equivalent of fungal vacuoles. We, therefore, looked for a vacuole phenotype in *fss1Δ* mutant spores using the vacuole lumen marker Cpy1-mCherry [49, 50]. The vacuoles of the diploid cell are excluded from developing spores and are formed de novo again in spores [51, 52]. This vacuole exclusion process appeared normal in asci produced by *fss1Δ* mutants (Figure S9A, see mCherry signal surrounding spores). The vacuoles formed within spores generated by *fss1Δ* homozygotes, however, showed mutant morphology. In wild-type, the Cpy1-mCherry signal was spread over several, small foci [53], whereas the spores produced by *fss1Δ* homozygotes generally had only one or two enlarged Cpy1-mCherry focus (Figure S9A, B). This vacuole fusion phenotype was dependent on the spore genotype as *fss1Δ/fss1+* heterozygotes showed a 2:2 distribution of those vacuole phenotypes (Figure S9A, B).

We next wanted to assay how this fused vacuole phenotype might affect the transmission of vacuoles to the daughter cells during the first cell division after germination. To do this, we stained the vacuoles in spores using the vacuole membrane dye, FM4-64, and imaged the distribution of the dye during the first cell division in germination. We saw that the total FM4-64 signal was the same in *fss1Δ* and wild-type spores, but the signal was restricted to a smaller area in the mutant spores, consistent with our observations with the Cpy1-mCherry marker (Figures 5A-C). As expected [54, 55], we found that more of the spore vacuole was generally retained in the mother cell during germination of spores generated by wild-type diploids. We previously observed that this asymmetry broke down in spores stressed by age or heat [55]. Similar to stressed spores, spores generated by *fss1Δ* homozygotes segregated spore vacuoles into the daughter cell more frequently than wild-type during germination (Figure 5C).

**Figure 5:**
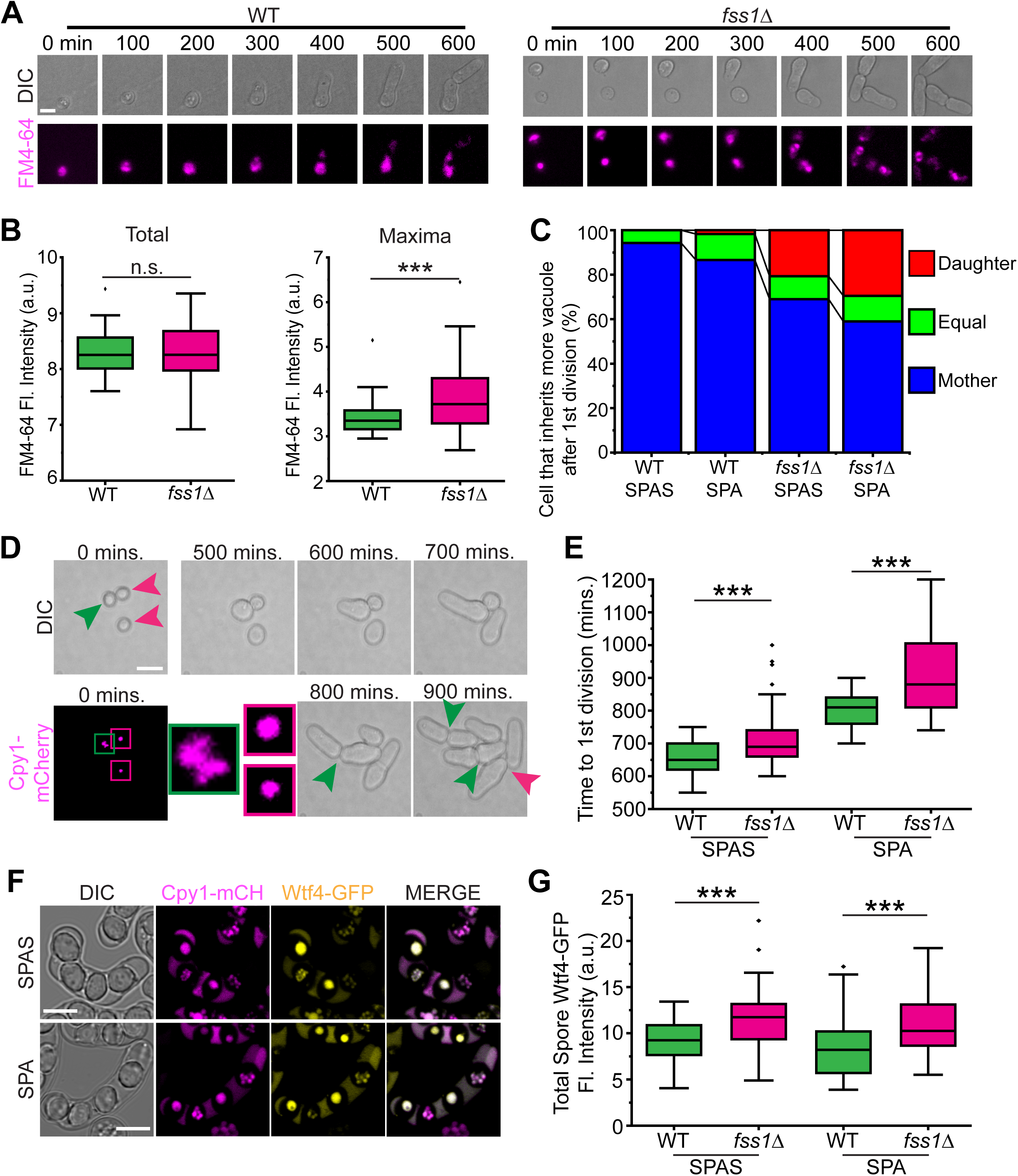
*fss1* expression in spores promotes proper vacuole formation and spore germination kinetics. A) Spores produced on SPAS from the indicated homozygous diploids were stained with FM4-64 to visualize the vacuoles and germinated on YEA+Sup. The cells were imaged every 10 minutes for 18 hours. Scale bar, 5 µm. B) The total FM4-64 fluorescence intensity (left) or the maximal FM4-64 fluorescence intensity (right) of the stained germinating spores described in A. n>50 in both samples; n.s.= not significant; ***, p<0.0001, t-test. C) The inheritance of FM4-64 stained vacuoles during germination of spores produced by homozygous diploids of the indicated genotypes on SPAS and SPA media. The mother cells were identified by the round shape of the spore that is retained during the first cell division. If the FM4-64 fluorescence intensity of the two cells post division was within 5% of each other, it was called equal between the mother and daughter. n>50 for all samples. D) Timelapse images of germinating spores derived from *fss1Δ* heterozygous diploids on SPAS or SPA plates. *cpy1-mCherry* was used to visualize the vacuoles. The spores were imaged on YES plates every 10 minutes for 24 hours. The green arrowheads indicate a wild-type spore (0 mins) and when it undergoes its first division (800 mins) and second divisions (900 mins). The magenta arrowheads indicate two *fss1Δ* spores (0 mins) and when one of the spores undergoes their first division (900 mins). The vacuole phenotype in the 0 timepoint was used to genotype the spores: wild-type spores have dispersed vacuoles (green box) and *fss1Δ* spores have fused vacuoles (magenta boxes). E) The time to the first division for wild-type and *fss1Δ* spores from the experiment shown in D. n>80 for all samples; ***, p<0.0001, t-test. F) Asci from *fss1Δ* heterozygous diploids after 48 hours on either SPAS or SPA mating media. The diploids were homozygous for *cpy1-mCherry*, to visualize the vacuoles and genotype *fss1*. The diploids were also homozygous for *wtf4-GFP*, which encodes two proteins that are degraded in spore vacuoles. The Wtf4-GFP clearance was used as a proxy for proper vacuole function in spores. Scale bars, 5 µm. G) Quantification of the fluorescence intensity of Wtf4-GFP in spores for the experiment shown in F. n>60 for all samples; ***, p<0.0001, t-test.

We used the clustered vacuole phenotype as a marker for spore genotype to compare germination dynamics of *fss1Δ* and *fss1*+ spores produced by heterozygotes. We found that both wild-type and *fss1Δ* spores germinated more slowly when they were generated on SPA, relative to the supplemented SPAS (Figure 5D, E). Importantly, we found that *fss1Δ* mutant spores germinate more slowly than their wild-type sister spores (Figure 5D, E; p<0.0001; t-test). This supports that both the vacuole and germination phenotypes are spore-autonomous, or dependent on the spore genotype, unlike the spore wall phenotypes described above.

The delayed germination phenotype we observe in *fss1Δ* mutants would put those spores at a competitive fitness disadvantage and likely contributed to why this gene was a strong hit in our TN-seq assay. It is possible the vacuole defect reflects a direct function of Fss1 in vacuole biology and that the delayed germination is due to the disrupted vacuole. It is also possible that the vacuole phenotype is a marker of cell stress and caused by some other spore-specific function of Fss1.

To test if vacuole function might be altered in *fss1Δ* spores, we assayed the degradation of tagged Wtf4 proteins. The *wtf4* is a member of a family of killer meiotic drivers that make two proteins that are destroyed in the vacuoles of spores [50]. We imaged asci produced by *fss1*+/ *fss1Δ* heterozygotes that were also homozygous for *cpy1-mCherry* and *wtf4-GFP*. We found that the GFP signal was higher in *fss1Δ* spores compared to their sister *fss1*+ spores (Figure F, G; p<0.0001: t-test). This is consistent with Fss1 acting to promote vacuole function (e.g. Wtf degradation) in spores.

## Discussion

### Novel roles in sexual reproduction for minimally characterized genes

*S. pombe* is a well-established model system with a relatively small genome (∼12.6 Mb) composed of a relatively small number of protein-coding genes (∼5000). Most of these genes have names and functional annotations through direct analysis in *S. pombe*, or have predicted functions due to varying degrees of homology with characterized genes in other organisms. Still, there are many genes with no names or known functions that have not been experimentally explored, beyond bulk studies. These largely uncharacterized genes include genes with human homologs and almost certainly includes genes with domains or molecular functions that are shared with human proteins.

In this work, we have provided an initial analysis of the role of 14 minimally characterized *S. pombe* genes in sexual reproduction (Table S1). These genes were identified as hits in a previous genome-wide saturation mutagenesis study that sought to quantify the contribution of all non-essential *S. pombe* genes to sexual reproduction. We found that all the tested genes did confer a defect in at least one assayed measure of sexual reproduction. Most of the phenotypes were, however, of mild to moderate severity. These subtle phenotypes, at least under standard laboratory conditions, likely contributed to these genes not being identified in previous studies. Still, it is important to note that genes do not have to have strong mutant phenotypes in our limited set of lab assays to be critically important for an organism’s fitness in the wild, as supported by the general conservation of the genes.

### Fss1 promotes spore wall development

We have assigned a name and function to a novel gene, *fss1*, a PQ-loop repeat family protein that has a unique dual role in spore development. First, Fss1 protein produced in a diploid cell undergoing meiosis facilitates spore wall development. One copy of the gene is sufficient for this role. In the complete absence of the protein, spore walls are thinner, with less chitin, and can have an altered smooth surface morphology.

PQ-loop proteins can act as tranceptors, with roles in both membrane transport and as membrane receptors [37, 56]. We propose Fss1 has specialized to focus on the later role as one of the critical prolines used in the conformational change that enables transport is changed to leucine [35]. This derived leucine is shared across fission yeast (Figure S3A), indicating the mutation has been conserved for over 200 million years [57]. We propose Fss1 acts as a receptor that facilitates spore wall development by promoting the recruitment of vesicles to the outer spore membrane (Figure 6A). We envision this role in directing vesicle fusion would be particularly important when resources are scarce (e.g. in auxotrophs or low nutrient media) and must be used efficiently to build strong spore walls.

**Figure 6:**
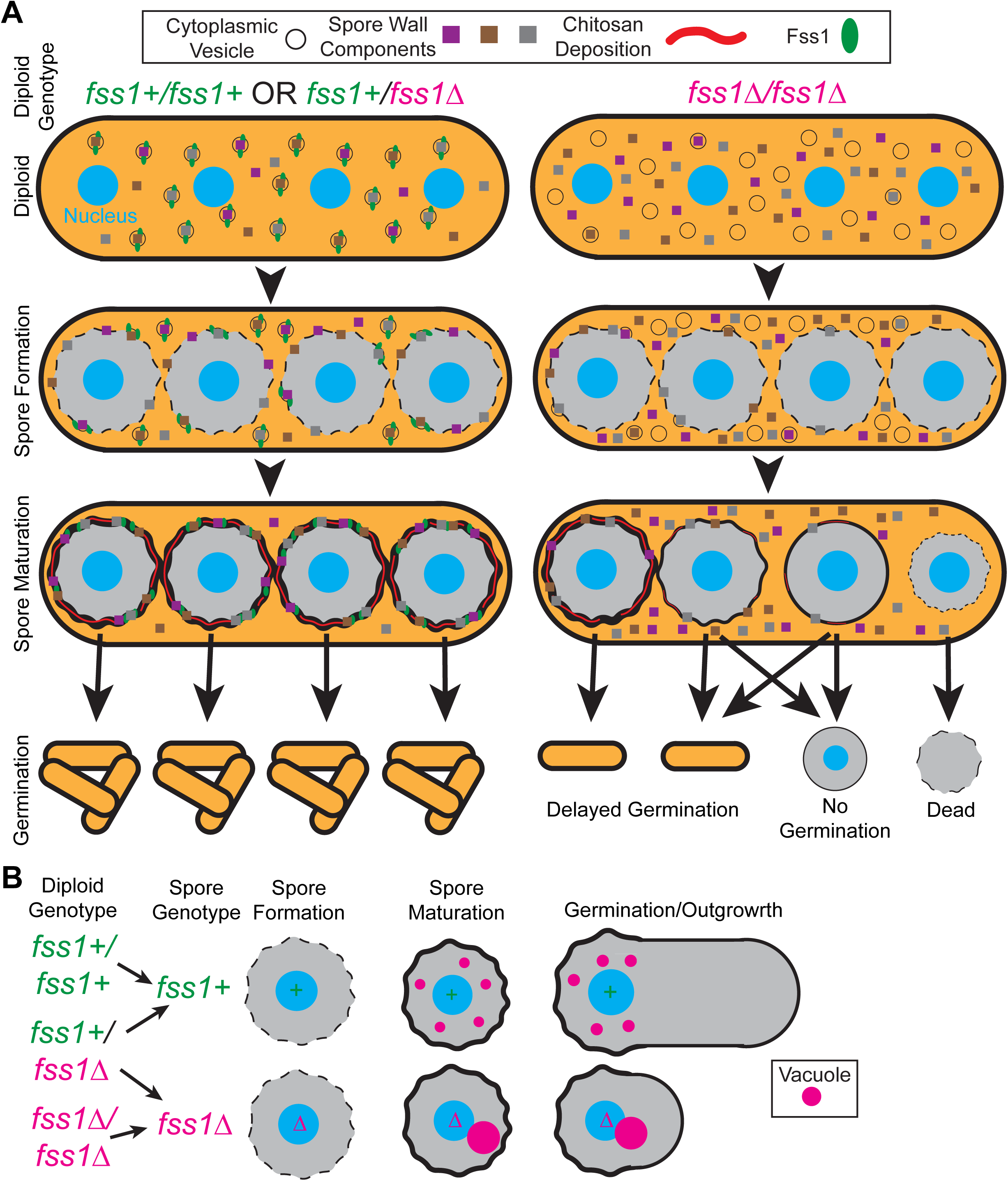
Model of Fss1 function in spore formation and germination. A) Model of Fss1 localization and function during spore formation. During meiosis, Fss1 localizes to cytoplasmic vesicles, but once the spore wall is initially formed, Fss1 is brought into the spore wall by merging cytoplasmic vesicles that help thicken and stabilize the spore wall. Fss1 may serve as a receptor for vesicle docking or additional protein recruitment as loss of *fss1* leads to an increase in spores whose walls are insufficient to survive spore maturation and/or germination. B) Model of spore-autonomous Fss1 function that promotes timely spore germination. In *fss1+* spores (top), the vacuoles are small, dispersed puncta and able to carry their functions. The spores germinate quickly and most of the vacuoles are retained in the mother cell. In *fss1Δ* spores (bottom), the vacuoles are aberrantly fused into one or two large foci. This affects the vacuole’s ability to clear deleterious proteins. The spores then germinate slower and vacuole inheritance loses its bias for the mother cell.

A previous study hypothesized that at least one spore-building protein (Mde10) was transported to the spore surface via fusion of vesicles to the outer forespore membrane [58]. We have now demonstrated this late vesicle fusion process occurs and that the vesicles can carry the Fss1 protein. We propose that Fss1-vesicles are delivering molecules and proteins important for spore wall deposition. These proteins could include Chs1, a transmembrane chitin synthase. *chs1* mutants have a similar phenotype to *fss1Δ*, wherein meiosis occurs normally, but spores fail to fully develop [59, 60]. *fss1*-facilitated vesicles could also be bringing in important spore maturation factors such as the GPI anchored protein Meu10 and/or the Mde10 transmembrane protein. These proteins share similar localization patterns as GFP-Fss1, in that they localize to the spore periphery during spore maturation [58, 61]. Mde10 is a particularly attractive candidate for Fss1-facilitated vesicle deliver as deletion mutants exhibit a loss of nubs along the spore wall, a phenotype observed in some spores produced by *fss1Δ* homozygotes (Figure 3C).

This model of spore building relies on delivery of factors from a shared pool produced in the diploid cell, so the genotype of the diploid, rather than the spore is important. The phenotypes of heterozygous mutants of the spore-building factors mentioned above have only been reported for *meu10* [61]. Like *fss1* heterozygotes, *meu10* heterozygotes build wild-type spore walls on all four spores, consistent with delivery of the protein to developing spores from a shared pool of protein from the diploid (Figure 6A). Also consistent with the model, *chs1* transcripts peak during meiosis [60]. Similarly, the Mde10 protein is produced primarily during meiosis, before spores individualize [58].

### Spore-expressed Fss1 affects vacuole morphology and promotes timely germination

Fss1 also has a second, distinct role within spores promoting timely germination and proper vacuole morphology, function and transmission during germination (Figure 5). Importantly, these vacuole and germination phenotypes depend on the genotype of the spore, which strongly implicates that additional *fss1* transcripts and proteins are produced after the spores individualize. We have not, however, identified the mechanism by which Fss1 acts in spores. Like our model for its role in spore formation, we speculate Fss1 could be acting as a receptor to direct proteins to the vacuole membrane whose loss could trigger vacuole fusion, such as Ypt4 [53, 62], Fab1 [63], Vps34 [64] or Ypt71 [65]. This would be analogous to the non-transport role of human PQLC2 in recruiting WDR41 to lysosomes [29]. Recently, loss of human PQLC2 also showed a loss of mTORC1 signaling and destabilization of the lysosome [66], thus Fss1 could be serving a similar role in vacuole stabilization. The slower germination speed of the *fss1Δ* spores could also be a result of the fused, inefficient vacuole; mislocalized vacuole proteins that affect germination, such as Ypt7 [52] and/or large vacuoles could simply be physically inhibiting the mitotic spindle and slowing cell division [63]. There is also the possibility that loss of Fss1 affects vacuole membrane composition, which in *S. cerevisiae,* can alter the vacuole morphology [67–69].

### Spore Maturation: A vital fusion step to a viable spore

While the gross mechanisms underlying forespore membrane formation have been described, the stage transitioning from forespore membrane completion to a mature, resistant spore (spore maturation) has not been studied as thoroughly. After the forespore membrane closes to cellularize the nascent spores, the spore wall must be constructed between the two layers of the of the forespore membrane to create the mature spore wall.

We propose an updated model for spore maturation in *S. pombe*. Initially, the forespore membrane encloses the nascent spore. At this stage, the bilayered forespore membrane contains little to no cell wall (Figure S10A, left column). Vesicles from the surrounding epiplasm (cytoplasm surrounding the spores) are then targeted to the new forespore membrane and merge with the outer layer to deposit cell wall components. This expands the luminal space between the spore membranes, but the burgeoning spore wall is still relatively smooth (Figure S10A, center column). When the spore wall has reached a point where it is very dense, then movement of additional deposited components becomes restricted, forming the more electrodense “nubs” (Figure S10A, right column). With the loss of *fss1*, this merging process of vesicles to the outer spore wall gets disrupted. Spore wall material fails to merge efficiently and can be displaced to the epiplasm. The spore wall is then thinner and more likely to fail to hold the spore cytoplasm as the spore matures (Figure S10B).

## Materials and Methods

### Strain Construction

We constructed the deletion strains using transformation cassettes amplified from the Pombe Deletion Collection (Bioneer) via PCR (Table S3). We introduced the deletion cassettes into SZY4976 (*h90, leu1-32, ura4-D18*) (Table S2) via a standard transformation approach [70] and verified via PCR (Table S3). For N-terminal GFP-tagging of Fss1, we used the *pFA6a-KanMX6-P41nmt1-GFP* plasmid (Addgene #39290) as the DNA template to generate a transformation cassette via PCR (Table S3). We verified the tagged gene by PCR and sequencing of the PCR product. We generated all other strains via crosses using standard approaches [71].

### Viable Spore Yield Assay

To assay the viable spore yield of *h90* strains, we inoculated single colonies in 5 mL of YEL and grew them to saturation overnight at 32°C. The next day, we plated 50 µL of the saturated culture to SPAS or SPA media in a single patch. At the same time, we diluted the saturated culture and plated these dilutions on YEA+Sup plates to determine the starting number of cells per µL. We grew these colonies on YEA+Sup for 3 days at 32°C, imaged and the colonies counted to approximate the number of cells plated to the SPAS or SPA plates. We incubated the mating patches on SPAS or SPA at 25°C for 7 days, after which the mating patches were scraped off and treated with glusulase and ethanol treatments to kill vegetative cells [72]. We then diluted the spores and plated the dilutions to YEA+Sup plates which were grown for 5 days at 32°C prior to counting the colonies to determine the number of viable spores present in the mating patch. We calculated the viable spore yield by dividing the number of viable spores present in the patches by the number of yeast cells originally plated [72]. All viable spores yield assays included a wild-type control, which was used to normalize the mean VSY of each experimental strain. We calculated the mean VSY for each strain from 6 biological replicates (6 single colonies).

To calculate VSY for experiments that included heterozygotes, we used *h+/h-* crosses. We first grew separate YEL cultures of the *h+* and *h-* strains. We then diluted the cultures to matching OD_600_ measured concentrations, mixed them and plated 50 µL of this combined culture onto the SPAS or SPA plate. We also used this combined culture to plate to the initial YEA+Sup plate.

For the *sdg1Δ* and *sdg2Δ* VSY assays, we used *S. pombe* conditioned YEA+Sup media as *sdg* deletions strains fail to grow at low concentration on YEA+Sup media [25]. We also grew the wild-type control strain on conditioned YEA+Sup media as well for normalization of the VSY.

### Mitotic Growth (Spot Dilution) Assays

We inoculated single colonies of each strain into 5 mL of YEL to grow to saturation overnight at 32°C. We then diluted each culture to an OD_600_ of 5 in 200 µL of YEL in a 96-well plate, generated ten fold serial dilutions, and spotted them to YEA+Sup plates (or *S. pombe* conditioned plates for *sdg* deletion strains) using a 48-well Multi-Blot Replicator (Frogger). We then incubated the plates at 30°C for 5 days and imaged them every day staring with the 3^rd^ day.

### Microscopy and Data Analysis

#### Isp3-GFP and mCherry-Psy1 fluorescence microscopy and tetrad imaging

We patched haploid *h90* cells to SPAS or SPA for ∼24 hours at 25°C. Then, we scraped a small portion of cells off the plate, resuspended them in 10 µL of water and then mounted 2.5 µL on microscope slide for imaging (Fisher Scientific, Cat#: 12-550-15). We imaged the cells using an Axio Observer.Z1 (Zeiss) wide-field microscope with a 63x Apochromat oil-immersion objective (1.4 NA). We took DIC images with a DIC II condenser and transmitted light. We acquired GFP images using a mercury lamp with a GFP filter cube that used a 440-470 nm bandpass filter, reflected off an FT 495 nm dichroic filter into the objective and emission collected using a 525-550 nm bandpass filter. We used a Hamamatsu ORCA Flash 4.0 camera using MicroManager software to acquire images.

#### Meiotic time-lapse imaging with *hht1-RFP*

We grew cells in 5 mL YEL liquid cultures overnight at 32°C to saturation. We then pelleted and washed the cells twice with water before resuspending them in 50 µL liquid SPAS media. We then plated 6 µL of this resuspension to a SPAS plate and then cut out a 0.6-cm-diamter punch of the agar which we then placed, cells-side down, into an 8-well µ-slide plate (Ibidi). We mounted the µ-slide plate in an environmental controlled chamber (Oko Lab) to keep the agar punches at a constant 25°C. We then imaged the cells with a Nikon Ti2 widefield microscope through a 60x Plan Apo (NA 1.4) oil objective. We imaged both transmitted light (Brightfield) and with a 561 nm laser (*hht1-RFP*). We collected a single Z-plane image every 10 minutes for 16 hours using a Prime 95B camera (Photometrics) using perfect focus. We analyzed the images using ImageJ software. We set karyogamy, as visualized by Hht1*-*RFP as t=0. We similarly followed the meiotic divisions using Hht1-RFP. We determined the time at which spores formed from brightfield images.

#### Spore germination time-lapse imaging

We prepared spores on SPA or SPAS using a standard approach, including glusulase and ethanol treatments [72]. We then plated 6 µL of the spore suspension to a YEA+Sup plate from which we cut a 0.6-cm-diamter punch of the agar. We then placed this agar punch, cells-side down, into an 8-well µ-slide plate (Ibidi) and obtained brightfield images from a single Z-plane every 10 minutes for 18 hours as described for the meiotic time course above, except the cells were incubated at 30°C. We analyzed the images in ImageJ.

To stain germinating spores with FM4-64, we first resuspended isolated spores in 3 mL of YEL and grew them at 30°C for 1.5 hours. We then added FM4-64 dye (Life Technologies) to a final concentration of 16 µM and incubated the cells for an additional 30 minutes at 30°C. We then pelleted and washed the cells twice with water and resuspended them in 1 mL of water. We plated this suspension onto YEA+Sup for imaging as described above, except a 561 nm laser was used to image the FM4-64 dye. We analyzed the images in ImageJ using hand-drawn regions of interest in which we measured integrated density for total fluorescence and maximum intensity (for the maxima measurement). In the frame where the first division after germination occurred, we measured the integrated density of the mother (which remains round at one end) and daughter cells. We considered the integrated density of the cells to be the same if the values were within 5% of each other (based on the higher integrated density value).

To analyze germination of spores generated by *fss1Δ* heterozygotes in Figure 5D-E, we prepared spores for germination as described above, but imaged them on a Nikon spinning disc microscope which consists of a Nikon Ti base coupled to a spinning disc head with a slide incubator set to 30°C (Yokogawa CSU-W1). We collected transmitted light and fluorescent images every 10 minutes for 24 hours and kept the cells in focus using Nikon Perfect Focus. We excited Cpy1-mCherry with a 561nm laser through a 100x Plan Apochromat Objective (Nikon, NA 1.45) and collected emission through a ET605/70 nm bandpass filter onto an sCMOS camera (Hamamatsu Flash 4). We analyzed the images in ImageJ, and determined the spore genotype by dispersed (wild-type) or fused (*fss1Δ*) *cpy1-mCherry* signal.

#### Cpy1-mCherry imaging of asci from heterozygous and homozygous crosses

We patched haploid *h+* and *h-* cells for heterozygous crosses (WT/*fss1Δ*) and haploid *h90* cells for homozygous crosses (WT/WT; *fss1Δ*/*fss1Δ)* to SPAS plates for ∼24 hours at 25°C. We then resuspended a small number of cells in 10 µL of water and imaged them on an Axio Observer Z1, LSM-980 (Zeiss) confocal microscope using a 63x Plan-Apochromat (NA 1.4) objective. We excited Cpy1-mCherry with a 561nm laser and its fluorescence was filtered through a 561-694nm bandpass filter with a pinhole set to 1 AU, onto a GaAsP-PMT. We also acquired transmitted light at the same time. We analyzed the images in ImageJ software. To determine the number of Cpy1-mCherry foci, we drew a region of interest around each spore and then used the number of maxima identified by the ‘find maxima’ process (prominence set to 1000) as the number of foci.

#### Spore wall staining and imaging

To stain the spore walls, we used *h90* wild-type strains with untagged *hht1* and *h90*, *fss1Δ::KanMX6, hht1-RFP* strains. For Figure S8C, D, we used haploid *h+* and *h-* cells for the heterozygous cross (WT/*fss1Δ*). We patched haploid cells to SPAS or SPA plates for 5 days at 25°C. We then scraped the patches off the plates into 500 µL of water. We then mixed equal concentrations of the untagged wild-type spores and *hht1-RFP, fss1Δ::KANMX6* spores in 20 µL of water. We then added either 1 µL of Calcofluor White (Sigma-Alrdich) or 1 µL of Aniline Blue (Sigma-Aldrich) dye to the spores, mixed and allowed the cells to sit for 15 minutes at room temperature. We then imaged the stained spores using an Axio Observer.Z1 (Zeiss) wide-field microscope with a 63x Apochromat oil-immersion objective (1.4 NA). We took DIC images with a DIC II condenser and transmitted light. We took RFP images using a mercury lamp with a DsRed filter cube that used a 530-585 nm bandpass filter, reflected off an FT 600 nm dichroic filter into the objective and emission collected using a 615 nm long-pass filter. We acquired Calcofluor White and Aniline Blue images using a mercury lamp with a DAPI filter cube that used a 365 nm green filter, reflected off an FT 395 nm dichroic filter into the objective and emission collected using a 445-450 nm bandpass filter. We took all images with a Hamamatsu ORCA Flash 4.0 camera using MicroManager software.

We analyzed all the images in ImageJ software by first drawing ROIs over each individual spore and then measuring the integrated density. We genotyped wild-type and *fss1Δ* spores using Hht1-RFP signal. Both dyes reduced the *hht1-RFP* signal about 3-fold compared to spores that were not dyed.

### Electron Microscopy and Data Analysis

#### Scanning Transmission Electron Microscopy (STEM)

We processed spores for STEM imaging as described in Nuckolls *et al.,* 2025 [55]. Briefly, we fixed the spores in 1.5% potassium permanganate (KMnO_4_) on ice for 30 minutes, then rinsed five times with ultrapure water for 10 minutes each. We then embedded the samples in 1% low-melt agarose [73], dehydrated through a graded ethanol series, and infiltrated and embedded in Hard Plus resin (Electron Microscopy Sciences). We microwaved (Pelco) the samples at 250 W for 3 minutes to assist resin infiltration at the 100% steps. We cut ultrathin sections (80 nm) using a Leica Actos ultramicrotome and collected the sections on slot grids. We post-stained the grids sequentially with 1% Sato’s Triple Lead for 3 minutes, 4% uranyl acetate in 70% methanol for 4 minutes, and again with 1% Sato’s Triple Lead for 5 minutes. We performed STEM imaging on a Zeiss Merlin SEM equipped with a STEM detector operated at 30 kV and 700 pA.

To determine spore wall thickness, we analyzed images with ImageJ software using hand-drawn ROIs for each spore. The first measurement was the entire area of the spore (wall and cytoplasm) and the second measurement was the area of the cytoplasm alone. Those areas were then assumed to be circles and converted into radii with the equation, A=πr^2^. The difference between the radius of the entire spore and radius of the cytoplasm was used as the spore wall thickness.

### Immuno STEM

We vacuum filtered yeast at OD 0.6 onto 0.45 µm filter paper (Millipore) subsequently kept moist on a media agar plate. We then loaded an aliquot into a 3mm x100 µm planchet (Technotrade) and fixed by high pressure freezing with a Leica ICE high pressure freezer. We immediately freeze substituted the samples using a Leica EM AFS2 system and fixed further in 0.2% uranyl acetate and 2% water in acetone and raised the temperature from −140°C to −40°C as follows: a four-hour ramp from −140°C to −90°C, an 8-hour hold, two 12 hour holds that took the samples to −80°C, then to −40°C where the samples were kept during washes. We first washed with acetone followed by two exchanges of ethanol over the span of 3 hours. We then infiltrated the samples with HM20 resin (Electron Microscopy Sciences) using the following dilutions at 2-3 hours each: 25%, 50%, 75% and then three exchanges at 100% with the final exchange sitting overnight. We then embedded the samples at −40°C and exposed them to UV light overnight with foil, then uncovered for 12 hours. We then warmed the samples from −40°C to 20°C overnight and ultimately left them to come to room temperature outside the FS unit. We then cut 60nm sections on nickel 300 mesh grids (Electron Microscopy Sciences) and blocked with 1% BSA + 0.1% glycine in PBST (0.1% Tween) for 1 hour. We then used a Rabbit anti-GFP antibody (gift from Mike Rout) at 1:50 and incubated for 1 hour. We washed the grids with PBST, then applied a colloidal gold conjugated goat, anti-rabbit secondary antibody (Jackson ImmunoResearch) at 1:50 for 1 hour. We then washed with PBST, then water before a 0.5% paraformaldehyde post fix for 30 minutes. We then washed the grids with water, dried and stained with 2% aqueous uranyl acetate for 2 minutes, dried and imaged on the same STEM system as described above.

#### Scanning Electron Microscopy (SEM)

We fixed spores in 50 mM sodium cacodylate buffer (pH 7.4) containing 2.5% glutaraldehyde, 2% paraformaldehyde, 1 mM CaCl₂, and 1% sucrose at 25°C for 2 hours, then stored the spores in fixative overnight at 4°C. We then processed the samples as described in Nuckolls *et al.*, 2025 [55] using the TOTO protocol [74]. Following dehydration, we critical-point dried the samples with a Tousimis Samdri-795. We mounted a section of Kimwipe onto an aluminum stub using carbon adhesive tape and sputter-coated (Leica ACE 600) the Kimwipe with 3 nm of Au/Pd. We then sprinkled spores onto the coated surface and then coated the entire stub with an additional 8 nm layer of carbon to reduce charging while minimizing surface texturing. We imaged at 2 kV and 200 pA using an InLens SE2 detector on a Zeiss Merlin scanning electron microscope.

We analyzed the SEM images using ImageJ software. We counted each spore in the field of view except spores that were out of the focal plane or shaded by the Kimwipe, as this could imitate “smoothness” on the outer spore wall. To be classified as a smooth spore, there had to be no difference in height of the spore wall along the entire visible spore.

#### Transmission Electron Microscopy (TEM)

We fixed yeast from SPAS media by high pressure freezing with a Leica ICE high pressure freezer. We then immediately freeze substituted the samples using a Leica EM AFS2 system and fixed with 2% osmium tetroxide, 2% uranyl acetate and 2% water in acetone brought to 0°C across an 86-hour temperature gradient. We then allowed the samples to come to room temp covered for an hour and then dehydrated the samples through 4, 1-hour incubations in acetone. We carried out resin infiltration into Hard Plus (Electron Microscopy Sciences) over six days with two resin exchanges per day running through the following percent dilutions with no accelerator in the resin: 10%, 20%, 30%, 50%, 70%, 90%, 100%, 100%. On the fifth day, we added the accelerator for 3 additional exchanges of 8-12 hours each and embedded the samples into small capsules in the evening on the sixth day. We polymerized the samples at 60°C for 48 hours, cut 60nm sections using a Leica Arctos ultramicrotome and post stained with 6 minutes each of 1% Sato’s Triple Lead and 4% uranyl acetate in 70% methanol. We imaged the grids on a ThermoFisher Talos Transmission electron microscope at 200kV.

---

**Table S1: List of 14 TN-Seq deletions that affect meiotic function.** The wild-type strain background (top) and the mutant alleles are listed with their systematic IDs, their given gene name in *S. pombe* (if applicable), and any predicted orthologous genes in humans (by HCOP) [26, 75]. The mating efficiency and sporulation efficiency were determined from DIC images taken 48 hours after plating haploids on the indicated medium (SPAS or SPA; see Figure S1B). Mating efficiency is the percentage of diploids and asci from total cells, sporulation efficiency is the percentage of asci from total diploids, atypical asci reflects the percentage of atypical asci from the total asci (see Figure S1C). The viable spore yield (VSY) of each deletion is the average of 6 independent experiments for each of two independent transformants (see Figure 1A, S1E) normalized to *leu1-32, ura4-D18* WT VSY. The spore germination rates were determined from DIC images of spores on YEA+Sup taken every 10 minutes for 18 hours at 32°C (see Figure S1D). The images were scored computationally using an approach that follows the loss of circularity as spores germinate [55]. The average time to the first division was also calculated computationally as the point when a single germinating spore could be delineated as two separate cells. The mitotic growth of the mutants was determined from spot assays of the cells on YEA+Sup (Figure S1A). Entries that are significantly different from the wild-type strain, without multiple test corrections, are shown with *p<0.05; **p<0.001, determined by t-test with wild-type strain.

**Table S2: Strain Table.** List of all the *S. pombe* strains used in the paper with their genotype and use in each figure.

**Table S3: Oligo Table.** List of all oligos used to make the 14 deletions and verify their deletions, as well as oligos used to construct and verify tagging Fss1 with GFP on the N-terminus. Oligos are listed by number, name, description of use and then sequence.

## Data Availabilty

All relevant data are within the manuscript and Supporting Information Files. Additional raw data and images can be found at the Stowers Original Data Repository at http://www.stowers.org/research/publications/LIBPB-2656. Instructions for accessing data from the Stowers Original Data Repository via FTP can be found at https://www.stowers.org/research/publications/odr#ftp.

## Acknowledgments

We thank the Zanders Lab and Core facilities at the Stowers Institute for help and guidance during the development and completion of this project. We thank Mike Rout for reagents used in this project. A.J.B. thanks V and V for support throughout the entire project.

---

**Figure S1:**
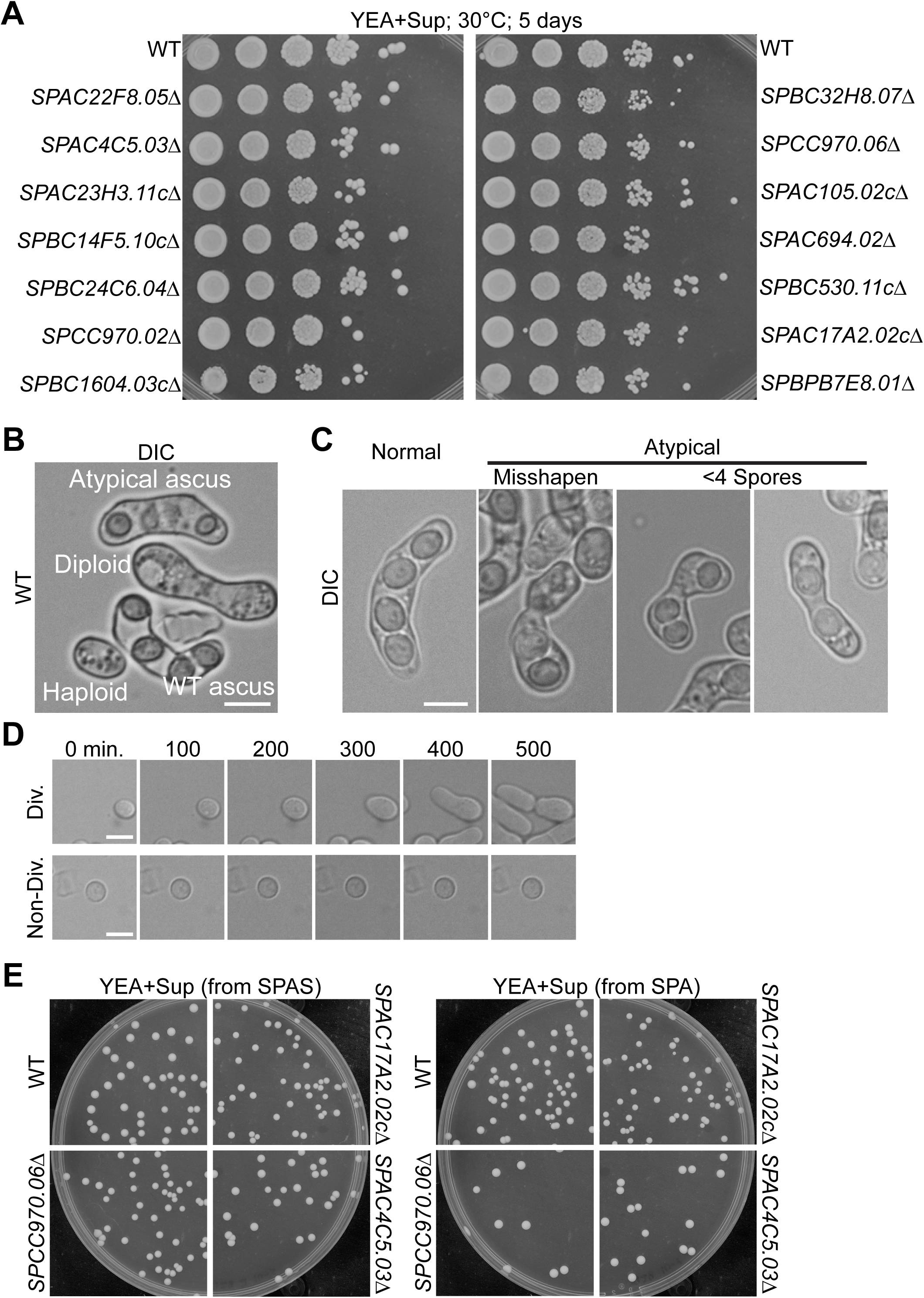
Analyses of mutants identified via TN-Seq. A) A tenfold serial dilution mitotic growth assay showing the growth of wild-type and all 14 tested mutants after 5 days on YEA+Sup at 30°C. B) Example image of an *h^90^* wild-type strain showing the four possible cell categories after 24 hours on sporulation medium (SPAS). Scale bar, 5 µm. C) Example images of a normal ascus (left) and atypical asci (right 3 panels) used in the quantification shown in Table S1, Figure 1E, and S6D. Scale bar, 5 µm. D) Timelapse images of wild-type spores germinating on YES plates. Spores were imaged every 10 minutes for 18 hours at 30°C. The top row shows a spore that germinates and divides. The bottom row shows a spore that failed to germinate within 18 hours. Scale bar, 5 µm. E) Example images of the post-meiosis (see Figure 1A) YEA+Sup plates (see Table S1). The images in the left column come from spores formed on SPAS media, and the images in the right column come from spores formed on SPA media.

**Figure S2:**
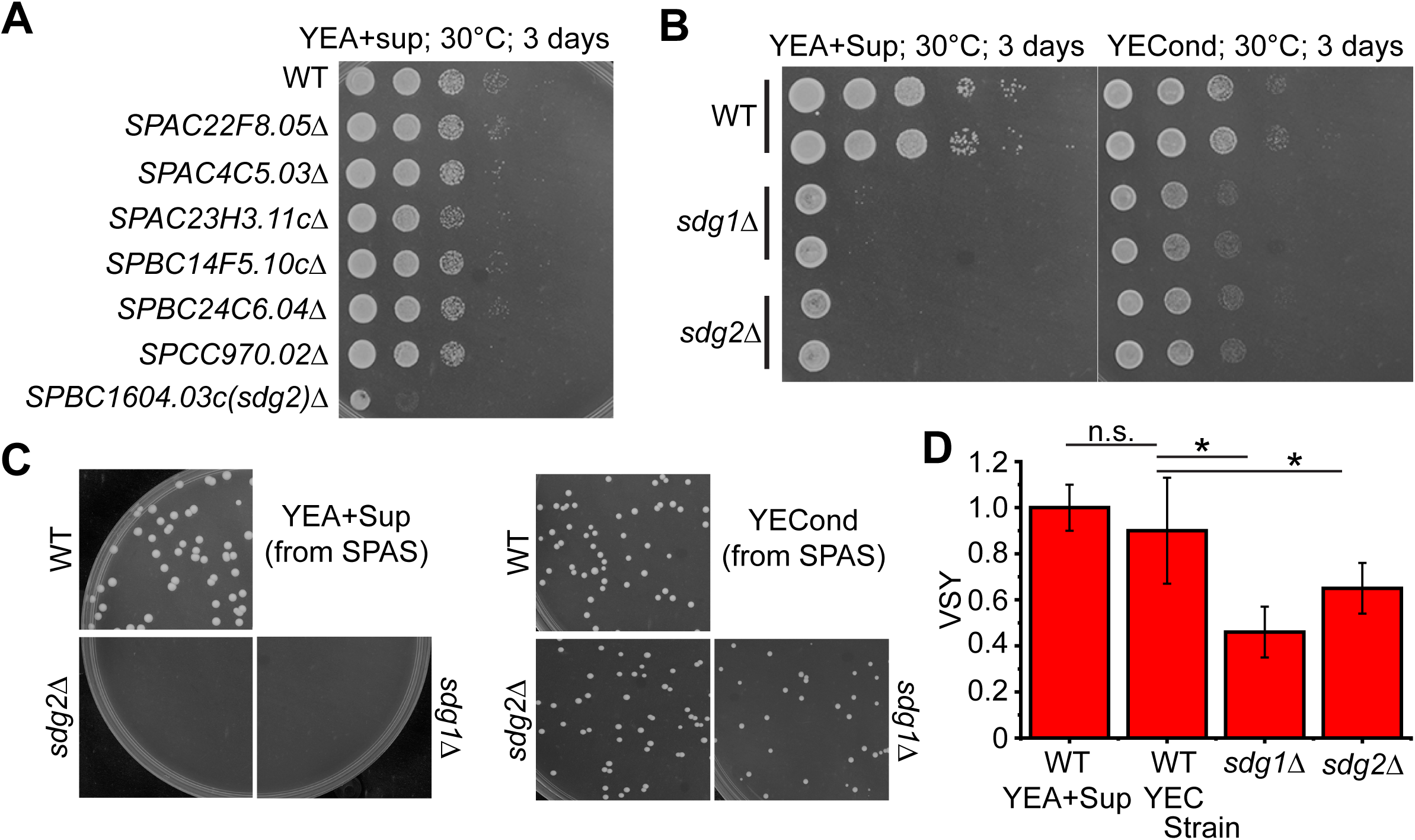
*SPBC1604.03c* is a Social Distancing Gene (*sdg)* that we name *sdg2*. A) Serial dilution mitotic growth assay of wild-type and seven mutants (including *SPBC1604.03c*) grown on YEA+Sup at 30°C after 3 days. B) Serial dilution mitotic growth assay of wild-type, *sdg1*Δ and *sdg2*Δ on YEA+Sup (left) or conditioned YE media (YECond or YEC) (right) at 30°C after 3 days. C) Example images of the post-meiosis (see Figure 1A) YES plates of wild-type, *sdg1Δ* and *sdg2Δ* strains. All spores were created on SPAS and germinated on YES (left) or conditioned YE media (right) at the same dilutions. D) Quantification of the viable spore yield (VSY) of wild-type, *sdg1Δ* and *sdg2Δ* strains on either YEA+Sup (wild-type only) or conditioned YE (YEC) plates (all strains). Each VSY is an average of 6 independent experiments and normalized to wild-type on YEA+Sup plates; n.s.= not significant; *p<0.05; t-test.

**Figure S3:**
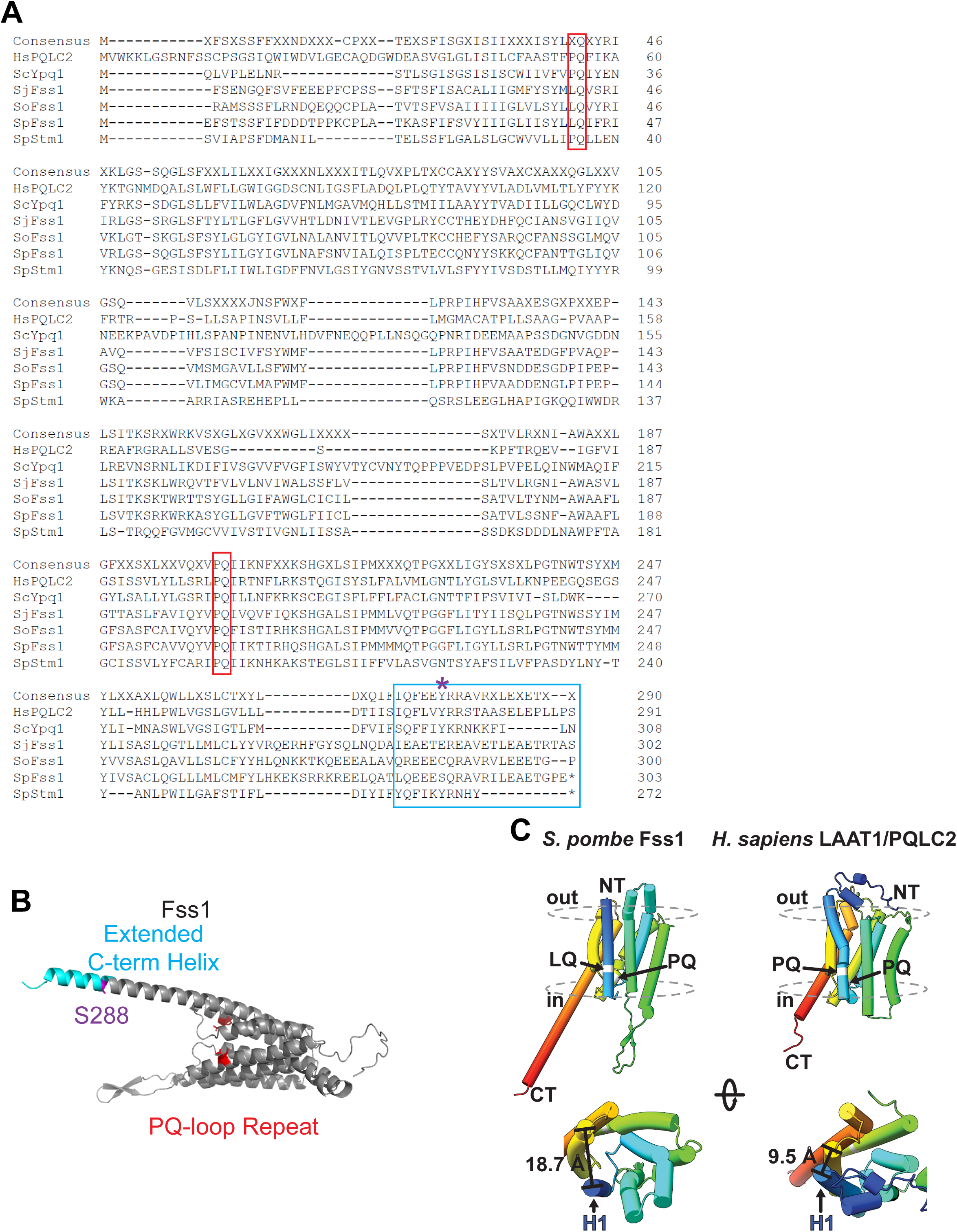
Fss1 is a PQ-loop repeat family protein that has retained a mutated PQ-loop for over 200 million years. A) Amino acid alignment of selected PQ-loop repeat family of proteins: *S. pombe* Fss1, *H. sapiens* PQLC2, *S. cerevisiae* Ypq1, *S. pombe* Stm1, *S. japonicus* Fss1 and *S. octosporus* Fss1. The red boxes denote the PQ-loop repeat regions of the proteins. The cyan box shows the extended C-terminus of Fss1. The purple asterisk above S288 indicates a possible TORC1 phosphorylation site identified in Berard et al. in *S. pombe* Fss1 [38]. B) AlphaFold prediction of Fss1 protein structure with PQ-loop repeat regions highlighted in red and extended C-term in cyan, which also contains possible phosphorylation residue, S288. C) Molecular model illustrating the PQ-loop repeats that function as hinges to facilitate amino acid transport in LAAT1/PQLC2. The mutation of the first PQ-loop to LQ, as in Fss1, is predicted to disrupt this hinge capability [35].

**Figure S4:**
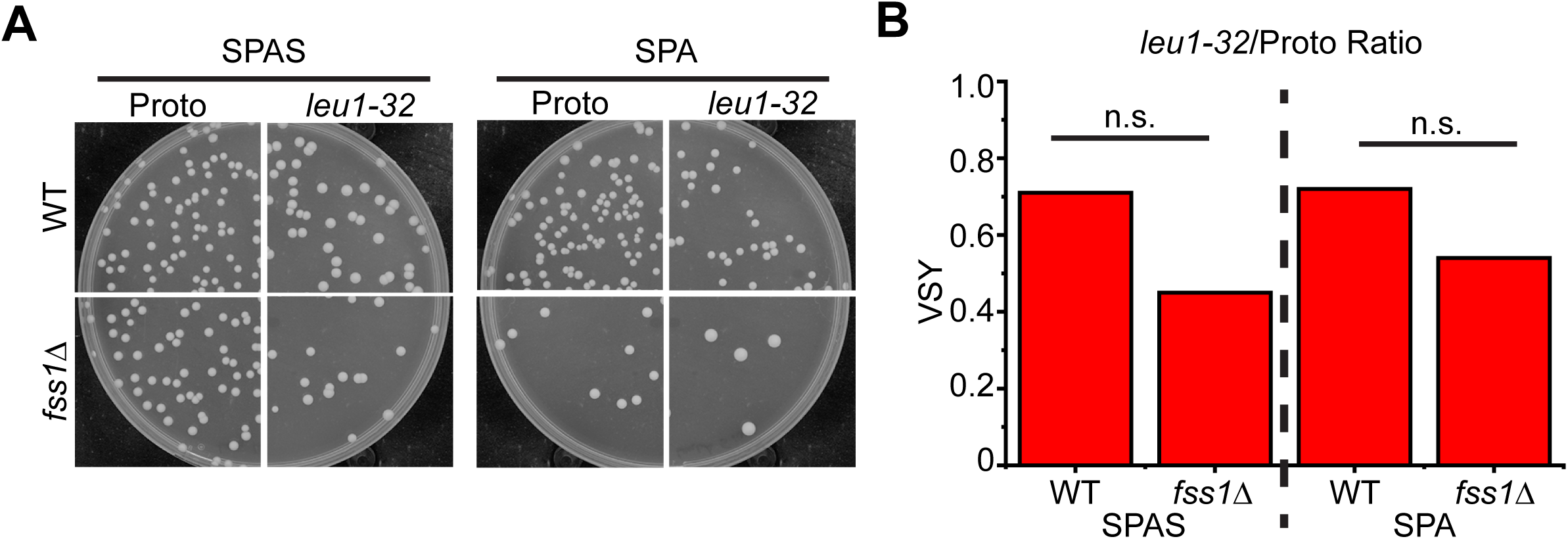
*fss1Δ* phenotype is not dependent on leucine auxotrophy. A) Example images of the post-meiosis (see Figure 1A) YEA+Sup plates from VSY assays from either prototrophic or leucine auxotrophic diploids of the given genotypes. The images in the left come from spores formed on SPAS media, and the right set of images come from spores formed on SPA media. The left column in each set is from prototrophic strains, while the right column in each set is auxotrophic for leucine (*leu1-32*). B) The ratio of the average auxotropic strain (*leu1-32*) VSY over the average prototrophic VSY for both mating media for wild-type and *fss1Δ* backgrounds. Each VSY was determined by 6 individual experiments for each strain; n.s. = not significant; chi-squared test.

**Figure S5:**
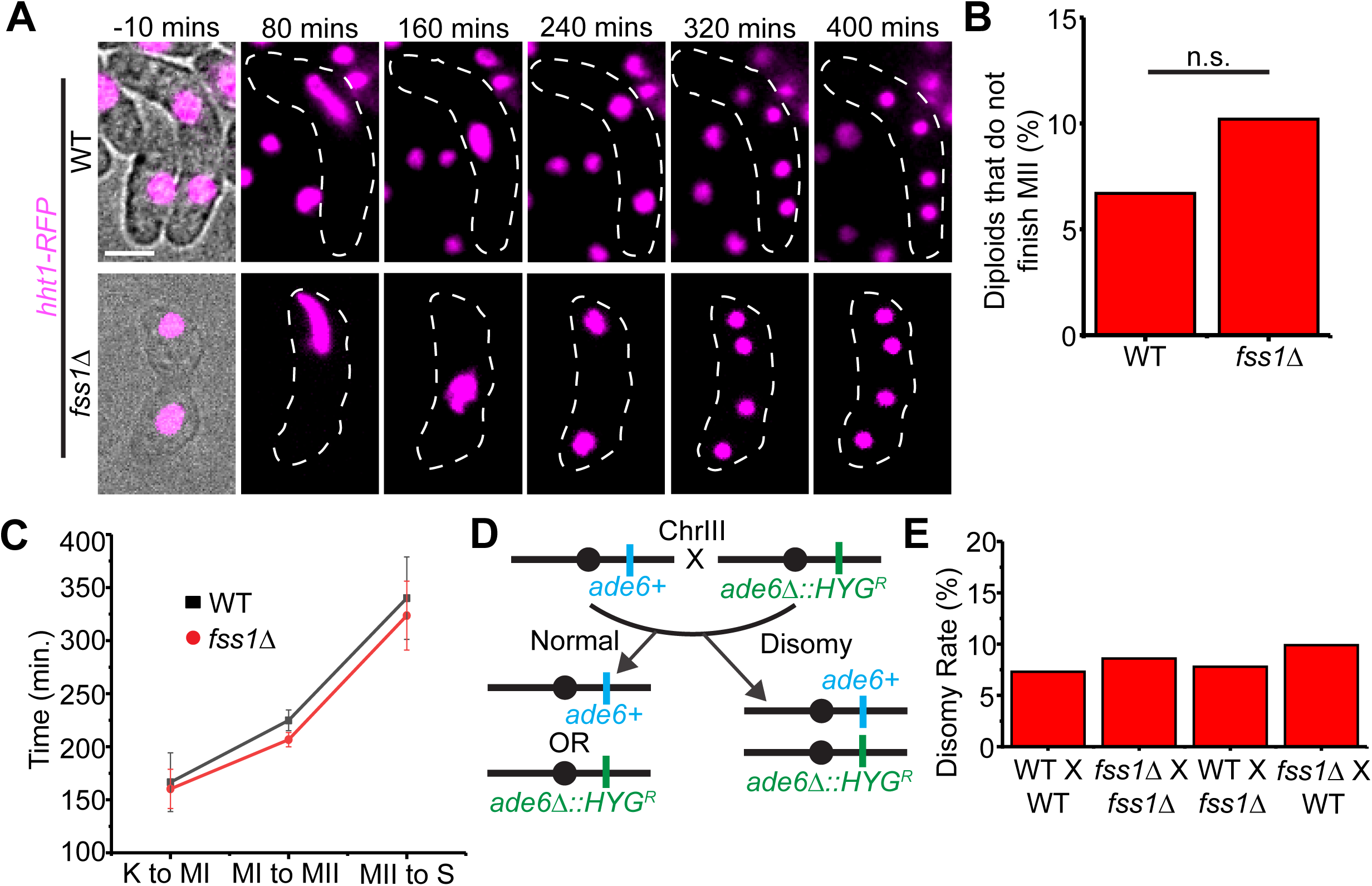
*fss1* is not required for meiotic chromosome segregation. A) Timeplase images of homozygous wild-type or *fss1Δ* diploids undergoing meiosis and sporulation on SPAS media. *hht1-RFP* labeled the nuclei and images were taken every 10 minutes for 16 hours. Karyogamy is the zero timepoint (t=0 mins, not shown). Scale bar, 5 µm. B) The percentage of diploids from A that failed to complete meiosis II (MII) during the 16 hours. n=90 (WT); n=88 (*fss1Δ)*, n.s. = not significant, chi-squared test. C) The cumulative time it took for wild-type and *fss1Δ* homozygous diploids (from A) to go through the meiotic divisions and sporulation. The zero timepoint (t=0) was karyogamy (K), and the steps measured were K to meiosis I (MI); MI to MII and MII to spore formation (S). n=45 (WT); n=48 (*fss1Δ*). D) Cartoon illustrating the chromosome III segregation assay. An *ade6+* haploid is mated with an *ade6Δ::HYG^R^*haploid. Their haploid progeny will inherit one of the markers. Chromosome missegregation in meiosis I can generate disomic spores carrying both markers. C) The fraction of viable spores that were chromosome III disomes recovered from the indicated crosses of wild-type and *fss1Δ* strains. The first listed strain was *h+, ade6+*, while second listed strain was *h-, ade6Δ::HYG^R^*. n=232 for all strains tested.

**Figure S6:**
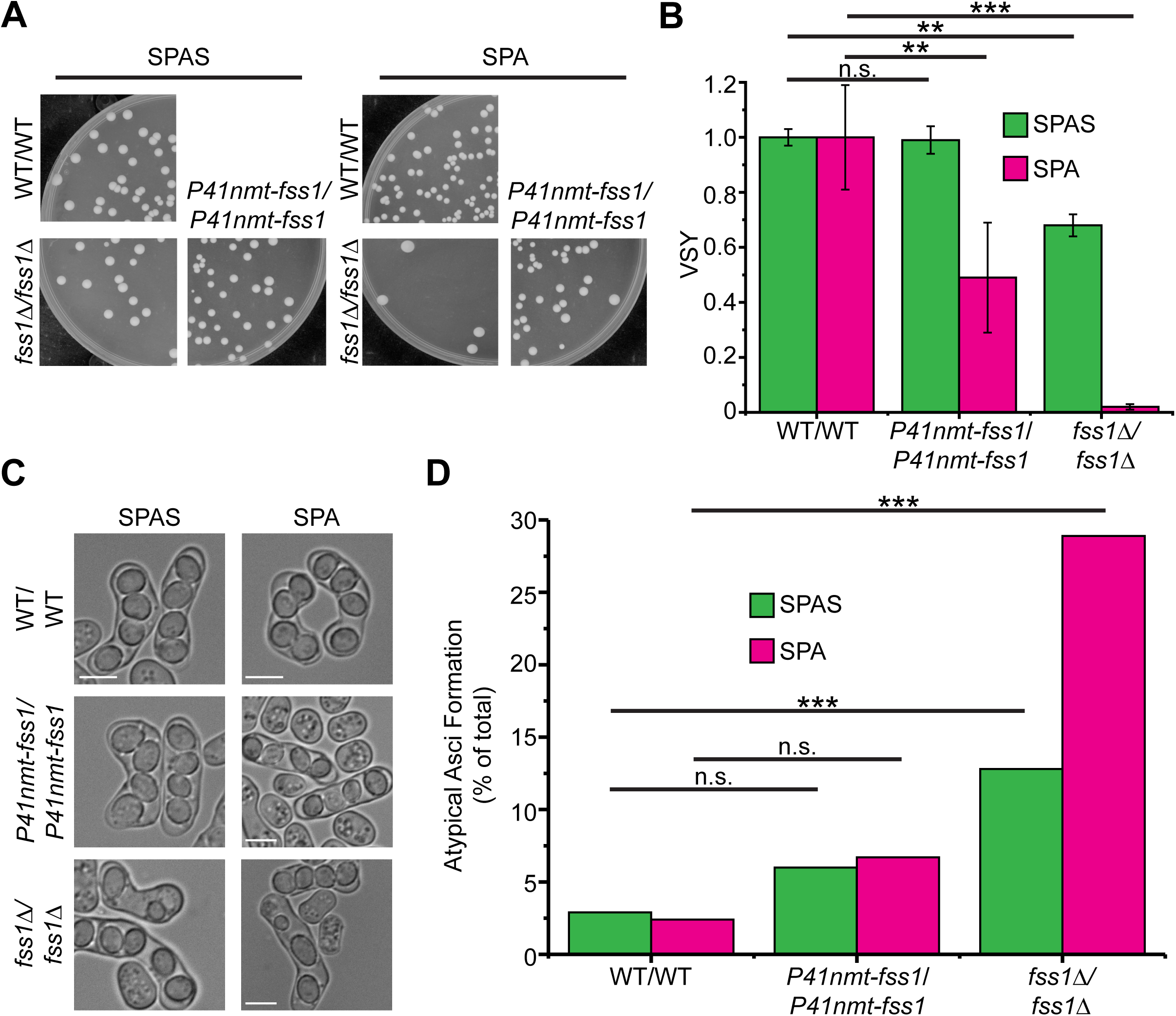
N-terminal tagging of Fss1 creates a mostly functional copy. A) Example images of the post-meiosis (see Figure 1A) YEA+Sup plates from the VSY assay. The images in the left set come from spores formed on SPAS, and the images in the right set come from spores formed on SPA. Diploids were either homozygous wild-type, homozygous *fss1Δ* or homozygous for the tagged *GFP*-*fss1* allele under the control of the *41nmt* promoter. B) Quantification of the average VSY for the indicated diploids on both mating media. Each average VSY was determined from 6 individual experiments. VSY was normalized to homozygous wild-type diploid for both mating media; n.s. = not significant; **p<0.001; ***p<0.0001; t-test. C) Example DIC images of asci derived from the indicated diploids after 48 hours on indicated media. The left column shows spores formed on SPAS, while the right column shows spores formed on SPA. D) Quantification of atypical asci quantified from the experiment shown in C (see Figure 1F, G; S1C). n>30 for all samples; n.s. = not significant; ***p<0.0001; chi-squared test.

**Figure S7:**
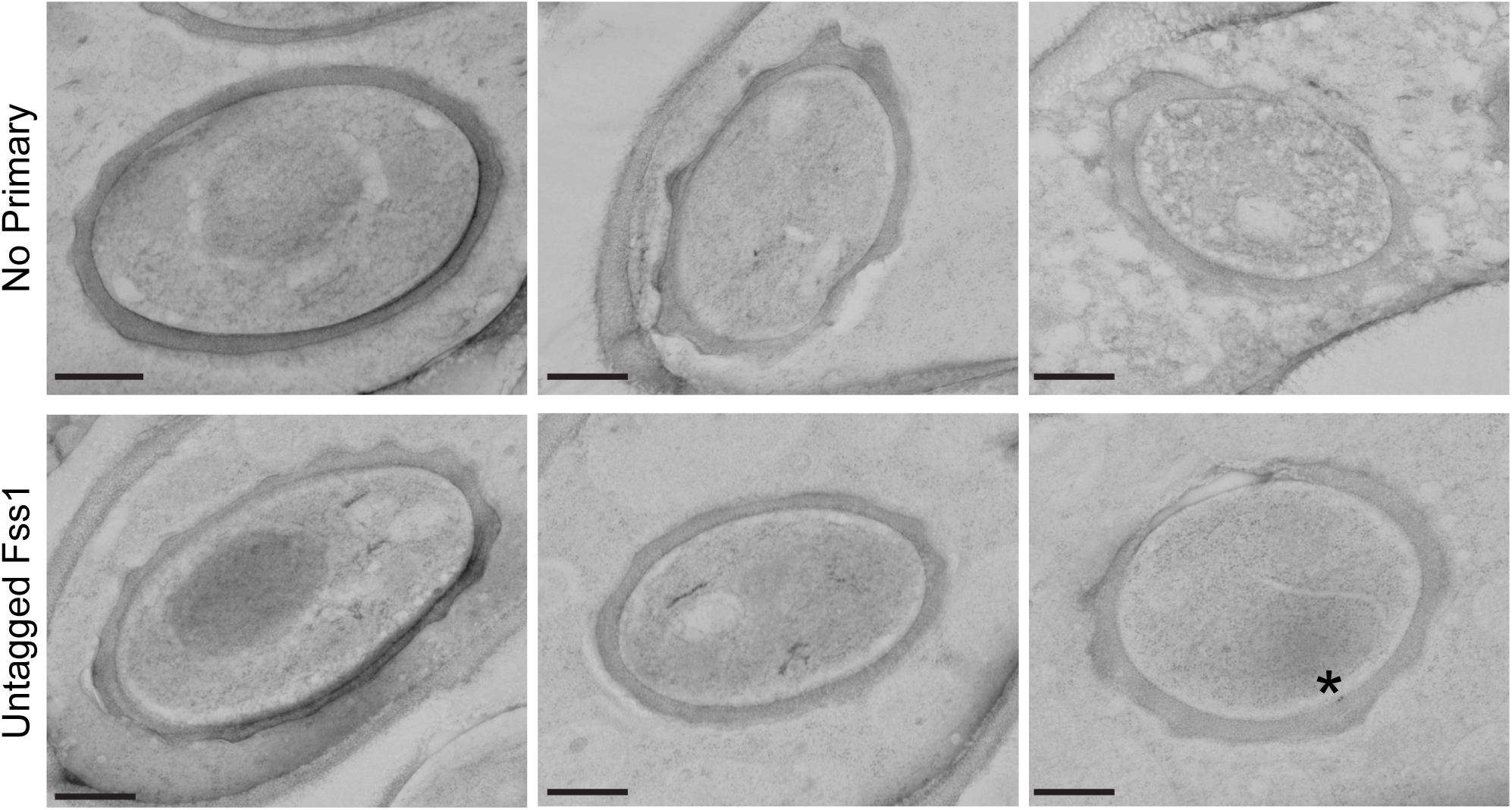
Controls for immunoEM show little to no non-specific labeling. Images of TEM slices during spore formation and maturation with controls for the GFP-Fss1 immunolabeling in Figure 2B. Top row, Spores with GFP-Fss1 present, but no primary antibody specific to the GFP has been added. Bottom row, Spores with both primary and secondary labeling, but the *fss1* gene was not tagged with GFP. There were a few instances of non-specific binding, but each contained only one or two particles localized to the inner spore membrane (marked with asterisk). Scale bars, 500 nm.

**Figure S8:**
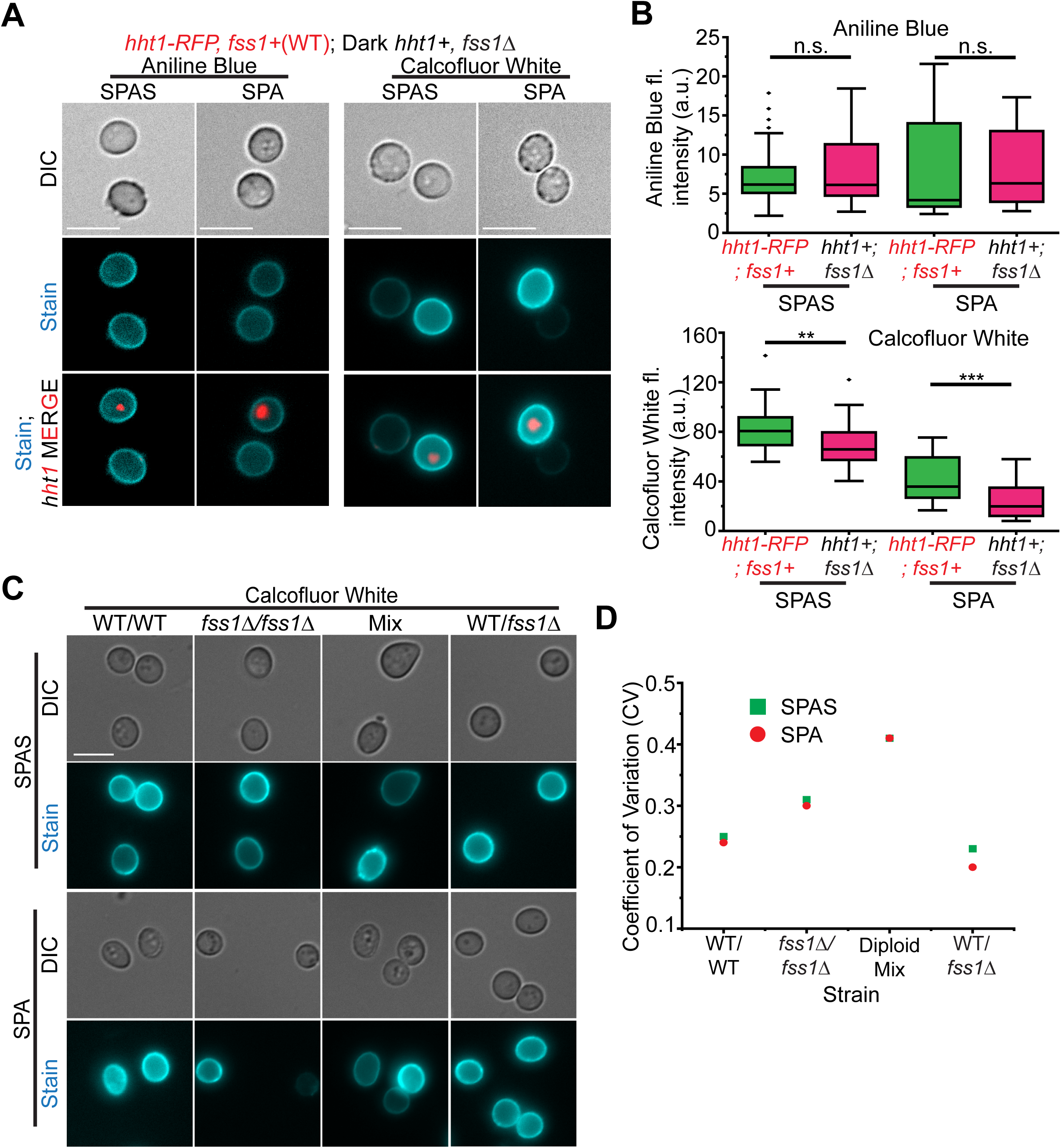
*fss1Δ* homozygous diploids show loss of chitosan (chitin) in spores. A) Images of spores derived from either wild-type diploids with tagged Hht1 (*hht1-mRFP)* or *fss1Δ* homozygous diploids (*hht1+*) (Reverse of the experiment in Figure 3E). The spores of distinct genotypes were created independently on either SPAS or SPA media, mixed together and stained with either Aniline Blue (left columns) or Calcofluor White (right columns). Scale bar, 5 µm. B) Quantification of Aniline Blue (top) or Calcofluor White (bottom) fluorescence intensity of spores from A. n>35 for all samples; n.s.= not significant; **p<0.001; ***p<0.0001; t-test. C) Images of spores stained with Calcofluor White made from either SPAS (top rows) or SPA (bottom rows). Spores were derived from wild-type diploids, *fss1Δ* homozygous diploids, a mixed population of the two homozygotes and *fss1Δ* heterozygous diploids. Scale bar, 5 µm. D) The coefficient of variation (CV; Standard deviation/Mean) of the Calcofluor White stained spores in C. n>90 for all samples.

**Figure S9:**
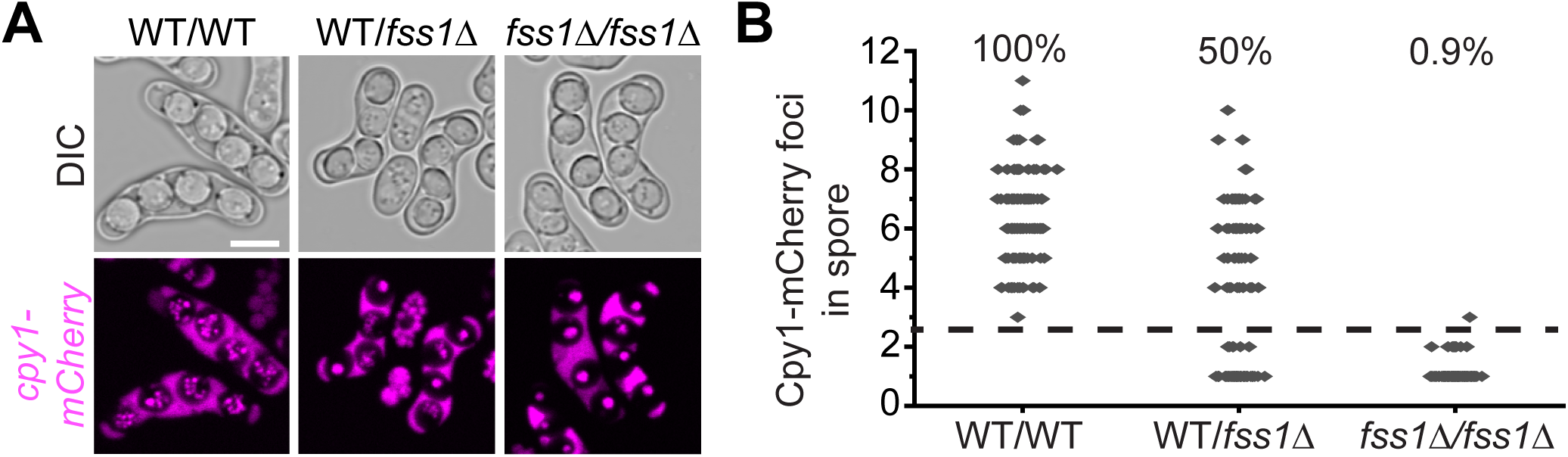
Loss of *fss1* expression in spores leads to abnormal vacuole fusion. A) Images of mature asci with vacuole marker, *cpy1-mCherry* formed by diploids of the given genotypes. The vacuoles found in the diploid parent cell are excluded from the spores and new vacuoles are made *de novo* in spores and are organized either as dispersed puncta (in WT spores) or as fused single foci (in *fss1Δ* spores). Scale bar, 5 µm. B) Dot plot showing the number of Cpy1-mCherry foci in each spore from the experiment shown in A. Dots above the dashed line had three or more *cpy1-mCherry* foci, while the dots below the dashed line had one or two foci. The indicated percentage at the top is the percentage of spores from each diploid that had three more or more foci. n>110 in all samples.

**Figure S10:**
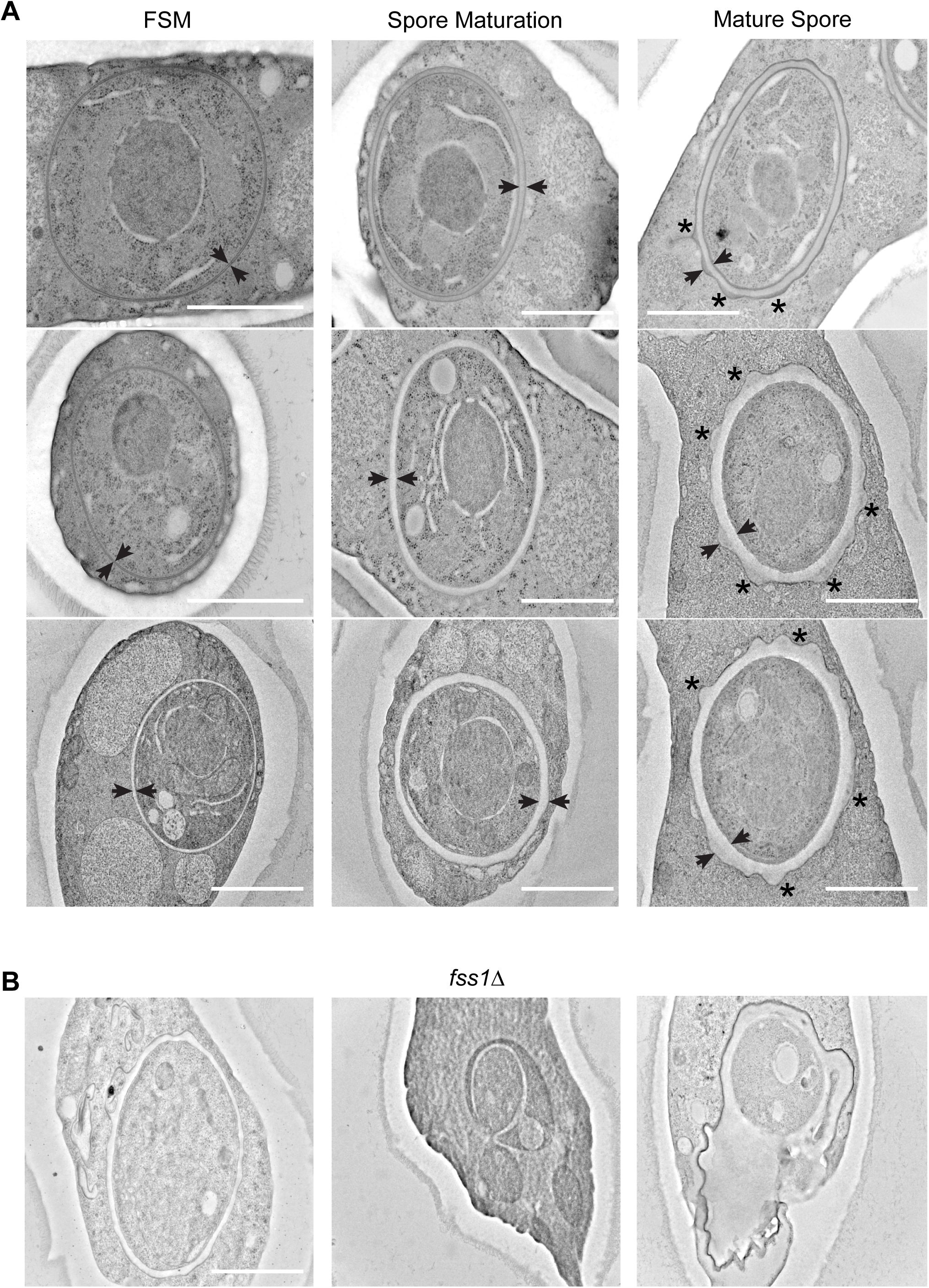
Spore wall maturation progresses through gradual expansion of the spore wall. A) TEM images of wild-type spores made on SPAS media. The forespore membrane (FSM) is a thin double bilayer with little to no luminal space (FSM; left column) that envelops each spore following meiosis. Our data (Figure 2) show that vesicles fuse to the complete FSM from the epiplasm as the spore wall thickens, while maintaining a smooth spherical shape (Spore Maturation; middle column). We propose that as the spore wall hardens, additional vesicles fuse and create the “nubs” around the spore wall (Mature Spore; right column). Arrowheads indicate the thickness of the FSM and/or spore wall and the distance between the two increases during maturation. Asterisks (*) indicate the “nubs” that have formed over the spore wall that are more electrodense than the neighboring spore wall underneath. Scale bar, 1 µm. B) TEM images of spores produced by *fss1Δ* homozygotes on SPAS media that created atypical spores. Spore wall material forms a membrane in the epiplasm away from the maturing spore wall (left image); spore wall rupture and leak of spore cytoplasm (center image) and/or inability to correctly fuse/compact the spore wall leading to “holes” in the spore wall (right image). Scale bar, 1 µm.

## References

1. Brawley, S.H. and L.E. Johnson, Gametogenesis, gametes and zygotes: an ecological perspective on sexual reproduction in the algae. British Phycological Journal, 1992. 27(3): p. 233–252.

2. Baker, B.S., et al., The genetic control of meiosis. Annu Rev Genet, 1976. 10: p. 53–134.

3. Goodenough, U. and J. Heitman, Origins of eukaryotic sexual reproduction. Cold Spring Harb Perspect Biol, 2014. 6(3).

4. Cohen, P.E. and J.W. Pollard, Regulation of meiotic recombination and prophase I progression in mammals. Bioessays, 2001. 23(11): p. 996–1009.

5. Martín-Castellanos, C., et al., A large-scale screen in S. pombe identifies seven novel genes required for critical meiotic events. Curr Biol, 2005. 15(22): p. 2056–62.

6. Blyth, J., et al., Genes Important for Schizosaccharomyces pombe meiosis identified through a functional genomics screen. Genetics, 2018. 208(2): p. 589–603.

7. Smolikov, S., K. Schild-Prüfert, and M.P. Colaiácovo, A yeast two-hybrid screen for SYP-3 interactors identifies SYP-4, a component required for synaptonemal complex assembly and chiasma formation in Caenorhabditis elegans meiosis. PLoS Genet, 2009. 5(10): p. e1000669.

8. Jain, D., et al., ketu *mutant mice uncover an essential meiotic function for the ancient RNA helicase YTHDC2*. Elife, 2018. 7.

9. Carrère, S., et al., A fully sequenced collection of homozygous EMS mutants for forward and reverse genetic screens in Arabidopsis thaliana. Plant J, 2024. 119(6): p. 3015–3026.

10. Sing, T.L., et al., Meiotic cDNA libraries reveal gene truncations and mitochondrial proteins important for competitive fitness in Saccharomyces cerevisiae. Genetics, 2022. 221(2).

11. Ucisik-Akkaya, E., J.K. Leatherwood, and A.M. Neiman, A genome-wide screen for sporulation-defective mutants in Schizosaccharomyces pombe. G3 (Bethesda), 2014. 4(6): p. 1173–82.

12. Yamashita, A., et al., Analysis of Schizosaccharomyces pombe meiosis. Cold Spring Harb Protoc, 2017. 2017(9): p. pdb.top079855.

13. Yamamoto, M., The molecular control mechanisms of meiosis in fission yeast. Trends Biochem Sci, 1996. 21(1): p. 18–22.

14. Dangarh, P., N. Pandey, and P.K. Vinod, Modeling the Control of Meiotic Cell Divisions: Entry, Progression, and Exit. Biophys J, 2020. 119(5): p. 1015–1024.

15. van Werven, F.J. and A. Amon, Regulation of entry into gametogenesis. Philos Trans R Soc Lond B Biol Sci, 2011. 366(1584): p. 3521–31.

16. Ohtsuka, H., et al., Sporulation: A response to starvation in the fission yeast Schizosaccharomyces pombe. Microbiologyopen, 2022. 11(3): p. e1303.

17. Shimoda, C., Forespore membrane assembly in yeast: coordinating SPBs and membrane trafficking. J Cell Sci, 2004. 117(Pt 3): p. 389–96.

18. Sakaguchi, T., et al., Diversity of spore morphology among fission yeasts. PLoS One, 2026. 21(8): p. e0355697.

19. Johnke, R. and G.M. Padilla, Germination and outgrowth of Schizosaccharomyces pombe spores isolated by a simple batch centrifugation technique. J Gen Microbiol, 1979. 115(1): p. 255–8.

20. Esposito, M.S. and R.E. Esposito, The genetic control of sporulation in Saccharomyces. I. The isolation of temperature-sensitive sporulation-deficient mutants. Genetics, 1969. 61(1): p. 79–89.

21. Esposito, R.E., et al., The genetic control of sporulation in Saccharomyces. II. Dominance and complementation of mutants of meiosis and spore formation. Mol Gen Genet, 1972. 114(3): p. 241–8.

22. Shimoda, C., et al., Characterization of meiosis-deficient mutants by electron microscopy and mapping of four essential genes in the fission yeast Schizosaccharomyces pombe. Mol Gen Genet, 1985. 200(2): p. 252–7.

23. Rabitsch, K.P., et al., A screen for genes required for meiosis and spore formation based on whole-genome expression. Curr Biol, 2001. 11(13): p. 1001–9.

24. Enyenihi, A.H. and W.S. Saunders, Large-scale functional genomic analysis of sporulation and meiosis in Saccharomyces cerevisiae. Genetics, 2003. 163(1): p. 47–54.

25. Billmyre, R.B., et al., Genome-wide quantification of contributions to sexual fitness identifies genes required for spore viability and health in fission yeast. PLoS Genet, 2022. 18(10): p. e1010462.

26. Carme, P., et al., PomBase in 202C: expanding knowledge, modeling connections. Genetics, 2026. 232(4).

27. Forsburg, S.L., The micromammals. G3 (Bethesda), 2024. 14(6).

28. Cui, Z., et al., Amino acid sensing and lysosomal signaling complexes. Curr Opin Struct Biol, 2023. 79: p. 102544.

29. Amick, J., et al., PǪLC2 recruits the CSorf72 complex to lysosomes in response to cationic amino acid starvation. J Cell Biol, 2020. 219(1).

30. Talaia, G., J. Amick, and S.M. Ferguson, Receptor-like role for PǪLC2 amino acid transporter in the lysosomal sensing of cationic amino acids. Proc Natl Acad Sci U S A, 2021. 118(8).

31. Chen, L.Ǫ., et al., A cascade of sequentially expressed sucrose transporters in the seed coat and endosperm provides nutrition for the Arabidopsis embryo. Plant Cell, 2015. 27(3): p. 607–19.

32. Guan, Y.F., et al., RUPTURED POLLEN GRAIN1, a member of the MtN3/saliva gene family, is crucial for exine pattern formation and cell integrity of microspores in Arabidopsis. Plant Physiol, 2008. 147(2): p. 852–63.

33. Kawano-Kawada, M., et al., Stm1 is a vacuolar PǪ-loop protein involved in the transport of basic amino acids in Schizosaccharomyces pombe. Biochim Biophys Acta Biomembr, 2021. 1863(2): p. 183507.

34. Chung, K.S., et al., Isolation of a novel gene from Schizosaccharomyces pombe: stm1+ encoding a seven-transmembrane loop protein that may couple with the heterotrimeric Galpha 2 protein, Gpa2. J Biol Chem, 2001. 276(43): p. 40190–201.

35. Liu, B., et al., LAAT-1 is the lysosomal lysine/arginine transporter that maintains amino acid homeostasis. Science, 2012. 337(6092): p. 351–4.

36. Ruivo, R., et al., Mechanism of proton/substrate coupling in the heptahelical lysosomal transporter cystinosin. Proc Natl Acad Sci U S A, 2012. 109(5): p. E210–7.

37. Saudek, V., *Cystinosin,* MPDU1, SWEETs and KDELR belong to a well-defined protein family with putative function of cargo receptors involved in vesicle trafficking. PLoS One, 2012. 7(2): p. e30876.

38. Bérard, M., L. Merlini, and S.G. Martin, Proteomic and phosphoproteomic analyses reveal that TORC1 is reactivated by pheromone signaling during sexual reproduction in fission yeast. PLoS Biol, 2024. 22(12): p. e3002963.

39. Billault-Chaumartin, I., et al., Actin assembly requirements of the formin Fus1 to build the fusion focus. J Cell Sci, 2022. 135(13).

40. Fukunishi, K., et al., The fission yeast spore is coated by a proteinaceous surface layer comprising mainly Isp3. Mol Biol Cell, 2014. 25(10): p. 1549–59.

41. Tanaka, K. and A. Hirata, Ascospore development in the fission yeasts Schizosaccharomyces pombe and S. japonicus. J Cell Sci, 1982. 56: p. 263–79.

42. Pérez, P., et al., Fission yeast cell wall biosynthesis and cell integrity signalling. Cell Surf, 2018. 4: p. 1–9.

43. Tahara, Y.O., M. Miyata, and T. Nakamura, Ǫuick-Freeze, Deep-Etch Electron Microscopy Reveals the Characteristic Architecture of the Fission Yeast Spore. J Fungi (Basel), 2020. 7(1).

44. Kippert, F. and D. Lloyd, The aniline blue ffuorochrome specifically stains the septum of both live and fixed Schizosaccharomyces pombe cells. FEMS Microbiol Lett, 1995. 132(3): p. 215–9.

45. Roncero, C., et al., Isolation and characterization of Saccharomyces cerevisiae mutants resistant to Calcoffuor white. J Bacteriol, 1988. 170(4): p. 1950–4.

46. Jézégou, A., et al., Heptahelical protein PǪLC2 is a lysosomal cationic amino acid exporter underlying the action of cysteamine in cystinosis therapy. Proc Natl Acad Sci U S A, 2012. 109(50): p. E3434–43.

47. Haq, M.R., et al., Immunolocalization of cystinosin, the protein defective in cystinosis. J Am Soc Nephrol, 2002. 13(8): p. 2046–51.

48. Town, M., et al., A novel gene encoding an integral membrane protein is mutated in nephropathic cystinosis. Nat Genet, 1998. 18(4): p. 319–24.

49. Sun, L.L., et al., Global analysis of fission yeast mating genes reveals new autophagy factors. PLoS Genet, 2013. 9(8): p. e1003715.

50. Nuckolls, N.L., et al., The wtf4 meiotic driver utilizes controlled protein aggregation to generate selective cell death. Elife, 2020. 9.

51. Roeder, A.D. and J.M. Shaw, Vacuole partitioning during meiotic division in yeast. Genetics, 1996. 144(2): p. 445–58.

52. Kashiwazaki, J., et al., A role for fission yeast Rab GTPase Ypt7p in sporulation. Cell Struct Funct, 2005. 30(2): p. 43–9.

53. Bone, N., et al., Regulated vacuole fusion and fission in Schizosaccharomyces pombe: an osmotic response dependent on MAP kinases. Curr Biol, 1998. 8(3): p. 135–44.

54. Plante, S., et al., Cell-surface copper transporters and superoxide dismutase 1 are essential for outgrowth during fungal spore germination. J Biol Chem, 2017. 292(28): p. 11896–11914.

55. Nuckolls, N.L., et al., Impacts of stress and aging on spore health in Schizosaccharomyces pombe. Microbiol Spectr, 2025. 13(12): p. e0171025.

56. Saudek, V., Potential dual function of PǪ-loop proteins such as cystinosin. J Biol Chem, 2017. 292(36): p. 15133.

57. Rhind, N., et al., Comparative functional genomics of the fission yeasts. Science, 2011. 332(6032): p. 930–6.

58. Nakamura, T., et al., ADAM family protein Mde10 is essential for development of spore envelopes in the fission yeast Schizosaccharomyces pombe. Eukaryot Cell, 2004. 3(1): p. 27–39.

59. Matsuo, Y., et al., Genetic analysis of chs1+ and chs2+ encoding chitin synthases from Schizosaccharomyces pombe. Biosci Biotechnol Biochem, 2004. 68(7): p. 1489–99.

60. Arellano, M., et al., Proper ascospore maturation requires the chs1+ chitin synthase gene in Schizosaccharomyces pombe. Mol Microbiol, 2000. 35(1): p. 79–89.

61. Tougan, T., et al., Meu10 is required for spore wall maturation in Schizosaccharomyces pombe. Genes Cells, 2002. 7(2): p. 217–31.

62. Rains, A., et al., Ypt4 *and lvs1 regulate vacuolar size and function in Schizosaccharomyces pombe*. Cell Logist, 2017. 7(3): p. e1335270.

63. Morishita, M. and C. Shimoda, Positioning of medial actin rings affected by eccentrically located nuclei in a fission yeast mutant having large vacuoles. FEMS Microbiol Lett, 2000. 188(1): p. 63–7.

64. Takegawa, K., et al., Vesicle-mediated protein transport pathways to the vacuole in Schizosaccharomyces pombe. Cell Struct Funct, 2003. 28(5): p. 399–417.

65. Kashiwazaki, J., et al., Two fission yeast rab7 homologs, ypt7 and ypt71, play antagonistic roles in the regulation of vacuolar morphology. Traffic, 2009. 10(7): p. 912–24.

66. Jeung, Y.J., et al., PǪ-loop repeat-containing 2 (PǪLC2) regulates mTORC1 lysosomal localization and autophagic ffux. Eur J Cell Biol, 2026. 105(3): p. 151544.

67. Leveille, C.L., et al., Yeast cells actively tune their membranes to phase separate at temperatures that scale with growth temperatures. Proc Natl Acad Sci U S A, 2022. 119(4).

68. Goodman, J.S., G.A. King, and E. Ünal, Cellular quality control during gametogenesis. Exp Cell Res, 2020. 396(1): p. 112247.

69. Sing, T.L., G.A. Brar, and E. Ünal, Gametogenesis: Exploring an Endogenous Rejuvenation Program to Understand Cellular Aging and Ǫuality Control. Annu Rev Genet, 2022. 56: p. 89–112.

70. Schiestl, R.H. and R.D. Gietz, High efficiency transformation of intact yeast cells using single stranded nucleic acids as a carrier. Curr Genet, 1989. 16(5-6): p. 339–46.

71. Forsburg, S.L. and N. Rhind, Basic methods for fission yeast. Yeast, 2006. 23(3): p. 173–83.

72. Smith, G.R., Genetic analysis of meiotic recombination in Schizosaccharomyces pombe. Methods Mol Biol, 2009. 557: p. 65–76.

73. Frankl, A., M. Mari, and F. Reggiori, Electron microscopy for ultrastructural analysis and protein localization in. Microb Cell, 2015. 2(11): p. 412–428.

74. Jongebloed, W.L., et al., Non-coating fixation techniques or redundancy of conductive coating, low kV FE-SEM operation and combined SEM/TEM of biological tissues. J Microsc, 1999. 193(Pt 2): p. 158–70.

75. Eyre, T.A., et al., HCOP: a searchable database of human orthology predictions. Brief Bioinform, 2007. 8(1): p. 2–5.

